# Cell-surface tethered promiscuous biotinylators enable small-scale surface proteomics of human exosomes

**DOI:** 10.1101/2021.09.22.461393

**Authors:** Lisa L. Kirkemo, Susanna K. Elledge, Jiuling Yang, James Byrnes, Jeff Glasgow, Robert Blelloch, James A. Wells

**Author notes:** **For correspondence:**, (James A. Wells). These authors contributed equally to this work (Lisa L. Kirkemo, Susanna K. Elledge).

## Abstract

Characterization of cell surface proteome differences between cancer and healthy cells is a valuable approach for the identification of novel diagnostic and therapeutic targets. However, selective sampling of surface proteins for proteomics requires large samples (>10e7 cells) and long labeling times. These limitations preclude analysis of material-limited biological samples or the capture of rapid surface proteomic changes. Here, we present two labeling approaches to tether exogenous peroxidases (APEX2 and HRP) directly to cells, enabling rapid, small-scale cell surface biotinylation without the need to engineer cells. We used a novel lipidated DNA-tethered APEX2 (DNA-APEX2), which upon addition to cells promoted cell agnostic membrane-proximal labeling. Alternatively, we employed horseradish peroxidase (HRP) fused to the glycan binding domain of wheat germ agglutinin (WGA-HRP). This approach yielded a rapid and commercially inexpensive means to directly label cells containing common N-Acetylglucosamine (GlcNAc) and sialic acid glycans on their surface. The facile WGA-HRP method permitted high surface coverage of cellular samples and enabled the first comparative surface proteome characterization of cells and cell-derived exosomes, leading to the robust quantification of 1,020 cell and exosome surface proteins. We identified a newly-recognized subset of exosome-enriched markers, as well as proteins that are uniquely upregulated on Myc oncogene-transformed prostate cancer exosomes. These two cell-tethered enzyme surface biotinylation approaches are highly advantageous for rapidly and directly labeling surface proteins across a range of material-limited sample types.

## Introduction

The cell surface proteome, termed the surfaceome, serves as the main communication hub between a cell and the extracellular environment (***Wollscheid et al., 2009***). As such, this cellular compartment often reveals the first signs of cellular distress and disease, and is of substantial interest to the medical community for diagnostic and therapeutic development (***Leth-Larsen et al., 2010***). The precise and comprehensive profiling of the surfaceome, termed surfaceomics, provides critical insights for our overall understanding of human health and can inform drug development efforts. Several strategies have emerged for either selective or comprehensive surfaceomics, including biocytin hydrazide labeling of surface glycoproteins (***Wollscheid et al., 2009***), chemical biotinylation of lysines via NHS-ester labeling (***Huang, 2012***), and promiscuous biotinylator fusion proteins (APEX2, BioID, SPPLAT) (***Rees et al., 2015a***; ***Sears et al., 2019***; ***Wollscheid et al., 2009***). While each of these strategies robustly label surface proteins, they: (1) require large sample inputs (biocytin hydrazide), (2) require production of genetically engineered cells (APEX2, BioID), (3) label only partner proteins by binding targeting antibodies fused to APEX2 or HRP (SPPLAT), (4) require extensive sample manipulation (biocytin hydrazide), or (4) exhibit increased nonspecific labeling (NHS-ester) (***Bausch-Fluck et al., 2012***; ***Elschenbroich et al., 2010***; ***Griffin and Schnitzer, 2011***; ***Kuhlmann et al., 2018***; ***Li et al., 2020b***). Moreover, many of these methods are not able to capture short and transient changes that occur at the cell surface, such as binding, adhesion, assembly, and signaling (***Kalxdorf et al., 2017***). These current methods complicate the direct characterization of small clinical samples such as extracellular vesicles in patient serum. As biological research increasingly depends on animal models and patient-derived samples, the requirement for simple and robust methods amenable to direct labelling of material-limited samples for proteomic analysis will become paramount.

**Table 1.**
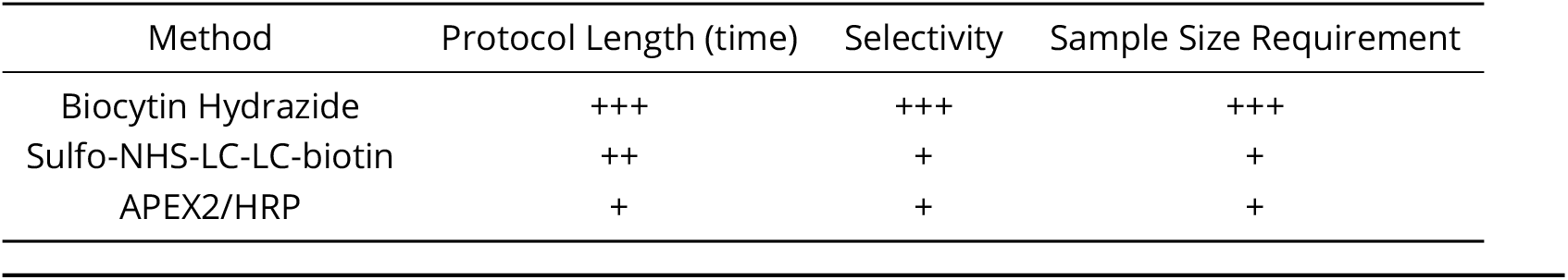
Current methods available for cell surface biotinylation.

| Method | Protocol Length (time) | Selectivity | Sample Size Requirement |
| --- | --- | --- | --- |
| Biocytin Hydrazide | +++ | +++ | +++ |
| Sulfo-NHS-LC-LC-biotin | ++ | + | + |
| APEX2/HRP | + | + | + |

Exosomes are small extracellular vesicles produced by both healthy and diseased cells (***Colombo et al., 2014***). In cancer, exosomes contribute to tumor growth and metastasis, modulate the immune response, and mediate treatment resistance (***Al-Nedawi et al., 2008***; ***Edgar, 2016***; ***Kalluri and LeBleu, 2020***; ***Shurtleff et al., 2018***). Consequently, these extracellular vesicles are a focus of intense clinical investigation. Recent studies suggest that exosomes incorporate proteins and RNA from the parent tumor from which they originate (***Lin et al., 2015***; ***Soung et al., 2017***), and certain proteins may be preferentially shuttled into exosomes (***Poggio et al., 2019***). There is also strong evidence that cancer-derived exosomes are unique from the exosomes derived from healthy surrounding tissues, and therefore represent a promising target for non-invasive, early-detection diagnostics or exosome-focused therapies (***Kalluri and LeBleu, 2020***; ***Skog et al., 2008***; ***Zhou et al., 2020***). However, strategies for the unbiased profiling of the exosomal membrane proteome remain limited. Isolation of high-quality, purified exosomes is challenging, requiring numerous centrifugation steps and a final sucrose gradient isolation, precluding the use of current labeling methods for membrane proteome characterization (***Poggio et al., 2019***; ***Shurtleff et al., 2018***). Strategies to characterize the exosome surface proteome would propel biomarker discovery and enable the differential characterization of the exosome proteome from that of the parent cell. These important studies could help illuminate mechanisms underlying preferential protein shuttling to exosomes.

Here, we functionalize the promiscuous biotinylators, APEX2 and HRP, as non-cellularly encoded exogenous membrane tethering reagents for small-scale surfaceomics, requiring <5e5 cells. This method is 10-100 fold more rapid than other existing protocols and requires fewer wash steps with less sample loss. Likewise, due to its selectivity towards tyrosines, it is not hindered by variability in individual protein glycosylation status (***Leth-Larsen et al., 2010***) or by impeding complete tryptic peptide cleavage through modification of lysines (***Hacker et al., 2017***), like biocytin hydrazide or biotin NHS methods, respectively. Using this robust new strategy, we performed surfaceomics on cells and corresponding exosomes from a cellular model of prostate cancer using the prostate epithelial cell line, RWPE-1 with or without oncogenic Myc induction. While certain proteins show increased expression in both parental cell and exosomal surfaces, a subset of proteins were found to be either pan-exosomal markers (MFGE8, IGSF8, and ITIH4) or selectively enriched with Myc overexpression in cancer-derived exosomes (ABCC1, SLC38A5, NT5E, FN1, and ANPEP). These differentially-regulated proteins pose interesting questions related to preferential protein shuttling, and the proteins upregulated in both cellular and exosomal contexts reveal candidates for early-stage urine or serum-based detection without invasive surgical intervention. We believe these simple, rapid, and direct labeling surfaceomic tools may be broadly applied to small-scale surfaceomics on primary tissues.

## Results

### Generation of promiscuous cell-surface tethered peroxidases for exogenous addition to cells

Both APEX2 and HRP are broadly used promiscuous proximity biotinylators that label nearby tyrosine residues in proteins through a radical intermediate mechanism using a biotin-tyramide reagent (**Figure 1A**) (***Hung et al., 2016***; ***Martell et al., 2016***). HRP has been targeted to specific cell-surface proteins through antibody conjugation to label target proteins and their binding partners (***Rees et al., 2015b***). More recently, HRP was used as a soluble cell surface labeler to identify rapid cell surface proteome changes in response to insulin (***Li et al., 2021***). Genetically encoded, membrane-targeted APEX2 and HRP have also permitted promiscuous labeling of proteins in specific cellular compartments, but these efforts required cellular engineering (***Hung et al., 2016***; ***Li et al., 2020a***). We sought to expand the use of these tools to biotinylate surface proteins of cells without the need for cellular engineering, enabling the specific enrichment of surface-resident proteins for mass spectrometry analysis.

**Figure 1.**
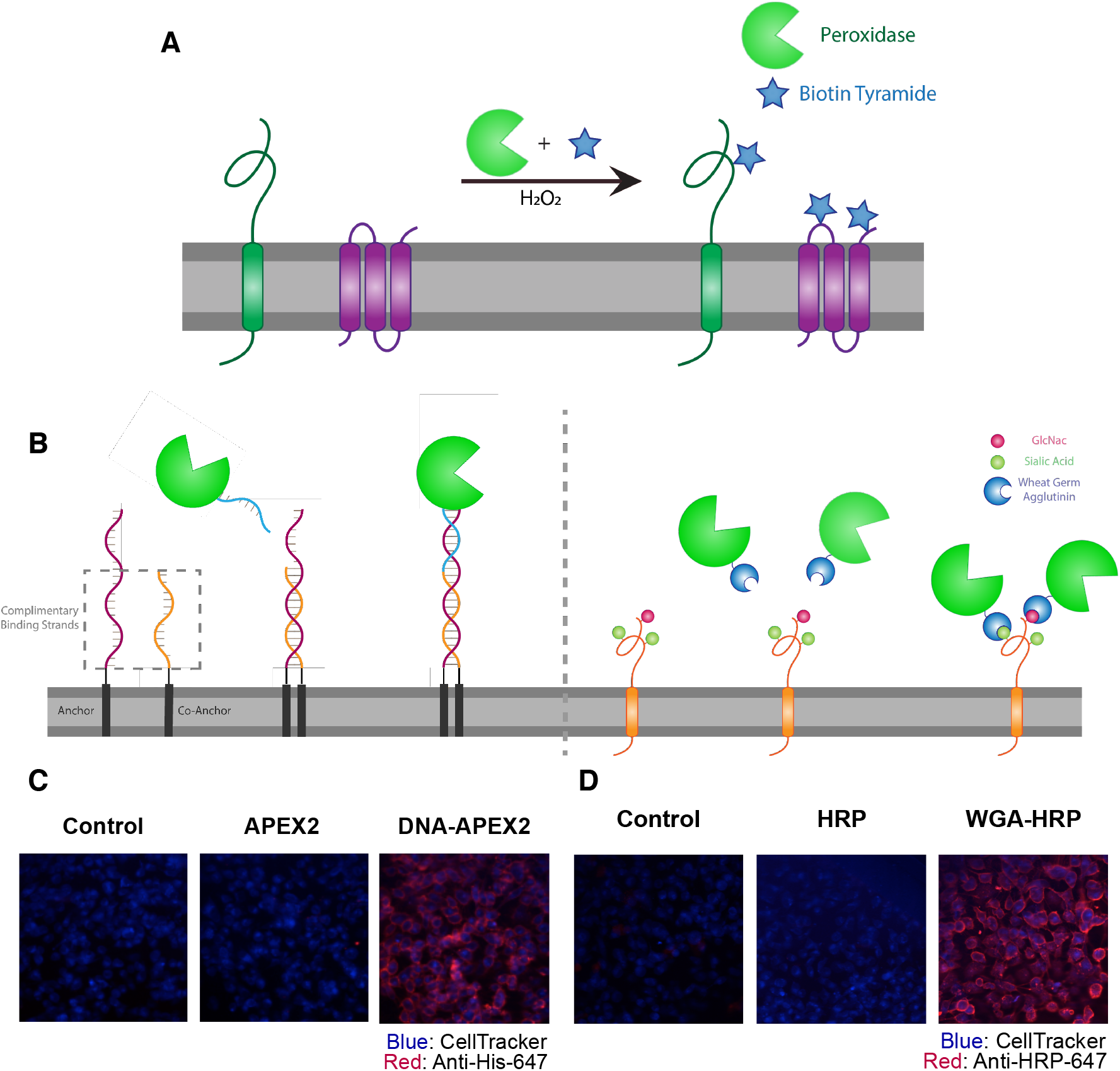
Direct labeling of promiscuous biotinylators to the cell membrane for rapid cell surface proteome characterization of small-scale biological samples. (A) Outline of enzymatic reaction mechanism. APEX2 and HRP both require biotin tyramide and hydrogen peroxide to produce the biotin-radical intermediate. (B) Tethering either enzyme is completed through differing mechanisms: (i) APEX2 is tethered through bio-conjugation of a single-strand of DNA, which is complementary to an exogenously added sequence of lipidated-DNA attached to the membrane, (ii) Commercially available wheat germ agglutinin-HRP associates with native GlcNAc and sialic acid glycan moieties on cell surface proteins. (C) Microscopy images depicting the localization of DNA-APEX2 to the cell surface of KP-4 cells after introduction of the lipidated-DNA complementary strands. (D) Microscopy images depicting the localization of WGA-HRP to the membrane of KP-4 cells.

The first approach we tested was to tether a DNA-APEX2 conjugate to the cell membrane through a lipidated DNA anchor. Gartner and co-workers have shown lipidated DNA anchors can tether together molecules or even cells (***McGinnis et al., 2019***; ***Weber et al., 2014***). Here the lipidated DNA is first added to cells, then hybridized with a complimentary strand of DNA conjugated to APEX2 (**Figure 1B**, left panel). To conjugate DNA to APEX2, we leveraged the single unpaired cysteine in the protein for site-specific bioconjugation of the complementary DNA. We first reacted APEX2 with DBCO-maleimide, after which the DBCO moiety was readily conjugated with azido-DNA. The kinetics of coupling was monitored using LC-MS and the conjugate was purified by nickel column chromatography, yielding a single conjugated product (**Figure 1 - Figure supplement 2A**) that retained full enzymatic function relative to unlabeled APEX2 (**Figure 1 - Figure supplement 2B**). Microscopy was used to observe the colocalization of DNA-conjugated APEX2 to the membrane (**Figure 1C**). This result was recapitulated using flow cytometry, indicating that this approach results in surface tethering of APEX2, an important step towards the specific labeling of the cell surfaceome (**Figure 1 - Figure supplement 2C**).

To avoid the need for bioconjugation, we also tested a commercially available reagent where the promiscuous biotinylator HRP is conjugated to the lectin wheat germ-agglutinin (WGA) (**Figure 1B**, right panel). WGA-HRP is used regularly in the glycobiology and neuroscience fields to label cell membranes for immuno-histochemistry and live cell imaging (***Mathiasen et al., 2017***; ***Wang and Miller, 2016***). This is an inexpensive and widely available tool that only requires the presence of surface protein N-acetylglucosamine (GlcNAc) and sialic acid glycans to localize HRP to the membrane. The successful colocalization of WGA-HRP to the plasma membrane compared to HRP alone was verified using immunocytochemistry, indicating this approach is a potential alternative for cell surface labeling (**Figure 1D**).

### Cell-tethered biotinylators more effectively label the surfaceome than non-tethered biotinylators and are comparable to biocytin hydrazide

Next, we set out to optimize labeling conditions for small-scale sample characterization. As APEX2 is kinetically slower than HRP (***Lam et al., 2015***), we used APEX2 to establish a suitable concentration range of enzyme for cell surface labeling. We found that 0.5 *μ*M APEX2 produced maximal labeling of cells (**Figure 2 - Figure supplement 1A**) and maintained equivalent labeling across a range of cell numbers (2.5e5 – 1e6 cells; **Figure 2 - Figure supplement 1B**). Next, we compared the efficiency of DNA-APEX2, WGA-HRP, and their non-tethered counterparts to biotinylate a small sample of 5e5 Expi293 cells. We found a 5- to 10-fold increase in biotin labeling for both tethered DNA-APEX2 and WGA-HRP relative to non-tethered controls as assessed by flow cytometry (**Figure 2A**) and western blotting (**Figure 2B**). Moreover, tethered DNA-APEX2 and WGA-HRP systems exhibited similar biotinylation effciency, suggesting either system is suitable for small-scale surfaceomics. Having both systems is useful, as some cells may not widely express glycoproteins recognized by commercially available lectin-HRP conjugates—such as some prokaryotic species—and therefore could require the glycan-agnostic DNA-tethered APEX2 construct (***Schäffer and Messner, 2017***).

**Figure 2.**
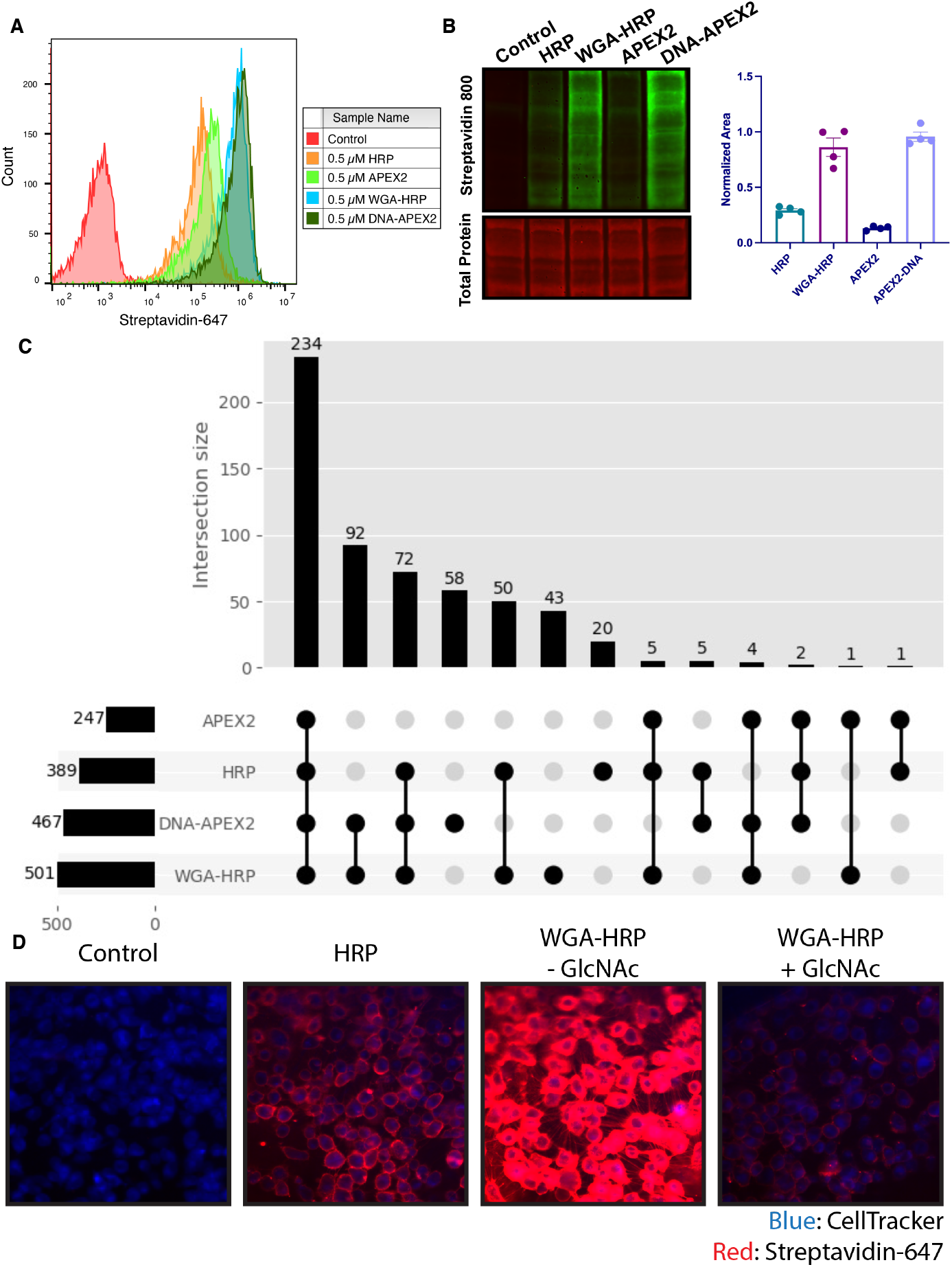
Membrane-localized peroxidases increases membrane proteome biotinylation compared to non-tethered counterparts. (A) Biotinylation of Expi293 cells treated with free enzyme (APEX2 or HRP) or cell-tethered enzyme (DNA-APEX2 or WGA-HRP) shown by flow cytometry. (B) Comparison of cell labeling with either free enzyme (APEX2 or HRP) or cell-tethered enzyme (DNA-APEX2 or WGA-HRP) shown by Streptavidin-800 western blot and total protein stain. Normalized area is plotted to the right. (C) Number of cell membrane proteins identified by mass spectrometry (>2 unique peptides, <1% FDR, found in all replicates) after treating 500,000 KP-4 pancreatic cancer cells with either free enzyme (APEX2 or HRP) or cell-tethered enzyme (DNA-APEX2 or WGA-HRP). (D) Microscopy images depicting extent of labeling with free HRP compared to WGA-HRP with and without the blocking sugar GlcNAc.

To compare the degree of surface protein enrichment these two systems offer, we enriched biotinylated proteins generated with either approach and compared the resulting enrichments using LC-MS/MS. As an initial efficacy comparison, cell surface labeling with DNA-labeled APEX2 or WGA-HRP was compared using 5e5 cells. In order to eliminate the possibility of suspension cell-specific results, we used a popular cell line model of pancreatic cancer, KP-4. We observed that the WGA-HRP identified slightly more plasma membrane annotated proteins (>2 unique peptides, found in all replicates) relative to DNA-APEX2, totaling 501 and 467, respectively. Notably, the number of IDs for both cell-tethered enzymes was higher than their untethered counterparts, with HRP identifying 389 cell surface proteins and APEX2 identifying 247 (**Figure 2C**). Importantly, in the upset plot shown, the group with the highest intersection includes all four enzyme contexts, showcasing the reproducibility of labeling through a similar free-radical based mechanism. The cell-tethered biotinylators also showed heightened surface enrichment compared to their untethered counterparts, as illustrated by the higher percentage of spectral counts derived from cell surface derived peptides (**Figure 2 - Figure supplement 2**). This suggests that localizing the enzyme to the membrane increases labeling of the membrane compartment, which we have previously observed with other enzymatic reactions (***Weeks et al., 2021***). As HRP is known to have faster kinetics compared to APEX2, it was unsurprising that WGA-HRP outperformed DNA-APEX2 in cell surface protein identifications. The heightened labeling of WGA-HRP was consistent with every cell line tested, including another pancreatic cancer model, PaTu8902, which resulted in an average of 848 cell surface proteins identified for WGA-HRP and 695 identified for DNA-APEX2 (**Figure 2 - Figure supplement 3**).

To confirm that the improved labeling by WGA-HRP was due to the binding of sugar units on the cell surface, we performed a sugar-blocking experiment with WGA-HRP using N-acetyl-D-glucosamine (GlcNAc) that would block the conjugate from binding to the cell. By pre-incubating WGA-HRP with excess N-acetyl-D-glucosamine, the ability of WGA-HRP to label the cell surface was markedly lower than WGA-HRP without GlcNAc as observed by microscopy (**Figure 2D**). A similar effect was also seen by flow cytometry (**Figure 2 - Figure supplement 4**). In addition, we also tested an on-plate protocol for simpler cell surface labeling of adherent KP-4 cells. We showed that cell surface labeling in this manner was comparable to labeling when the cells were in suspension (**Figure 2 - Figure supplement 5**).

As WGA-HRP consistently outperformed DNA-APEX2 by proteomics and represents a more facile method amenable to broad application in the field, we chose to compare the proteomic labeling results of WGA-HRP to other standard cell surface labeling methods (sulfo-NHS-LC-LC-biotin and biocytin hydrazide) on a prostate epithelial cell line, RWPE-1 with and without oncogenic c-Myc overexpression. Sulfo-NHS-LC-LC-biotin reacts with primary amines to form amide conjugates but has notoriously high background contamination with intracellular proteins (***Weekes et al., 2010***). Biocytin hydrazide labeling is a two-step process that first involves oxidizing vicinal diols on glycoproteins at the cell surface, then reacting the reactive aldehyde byproducts with biocytin hydrazide (***Elschenbroich et al., 2010***). Both WGA-HRP and biocytin hydrazide were able to identify similar numbers of cell surface proteins, with sulfo-NHS-LC-LC-biotin detecting the highest number of overall surface proteins. (**Figure 3 - Figure supplement 1A**) However, the cell surface enrichment levels were notably higher in both WGA-HRP and biocytin hydrazide (**Figure 3 - Figure supplement 1B**), suggesting a larger portion of the total sulfo-NHS-LC-LC-biotin protein identifications were of intracellular origin, despite the use of the cell-impermeable format. All three methods were highly reproducible across replicates (**Figure 3 - Figure supplement 2A-C**). Compared to existing methods, WGA-HRP not only labels cells efficiently with much lower input material requirements, it is also able to enrich for cell surface proteins to a similar extent in a fraction of the time.

### WGA-HRP identifies surface markers of Myc-driven prostate cancer in both cells and exosomes

Prostate cancer remains one of the most common epithelial cancers in the elderly male population, especially in Western nations (***Litwin and Tan, 2017***; ***Rawla, 2019***). While metastatic progression of prostate cancer has been linked to many somatic mutations and epigenetic alterations (PTEN, p53, Myc etc.), more recent work determined that alterations in Myc occurs in some of the earliest phases of disease, i.e. in tumor-initiating cells (***Koh et al., 2010***). This finding promotes the idea that the development of early-stage diagnostic tools that measure these Myc-driven disease manifestations could improve detection and overall patient disease outcomes (***Koh et al., 2010***; ***Rebello et al., 2017***). One mode of early detection that has gained prominence is the use of prostate cancer-derived exosomes in patient serum and urine (***Duijvesz et al., 2013***; ***McKiernan et al., 2016***). Exosomes are known to play important roles in the progression of prostate cancer, including increasing tumor progression, angiogenesis, metastasis, and immune evasion, making this subcellular particle an extremely informative prognostic tool for disease progression (***Akoto and Saini, 2021***; ***Lorenc et al., 2020***; ***Saber et al., 2020***).

To elucidate promising targets in Myc induced prostate cancer, we utilized our WGA-HRP method to biotinylate cells from both normal epithelial prostate cells (RWPE-1 EV) and oncogenic Myc-induced prostate cancer cells (RWPE-1 Myc, **Figure 3A**). Importantly, by using an isogenic system, we are able to delineate specific Myc-driven protein expression changes, which could be helpful in the identification of non-invasive, early-detection diagnostics for cancer driven by early Myc induction. In addition to having marked overexpression of c-Myc in the RWPE-1 Myc cells compared to the EV controls (**Figure 3B**), they also grow with a more mesenchymal and elongated morphology compared to their EV counterparts (**Figure 3C**), which would suggest large cell surface changes upon oncogenic Myc induction. We initially used WGA-HRP to quantitatively compare the cell surface profiles of Myc-induced prostate cancer to the EV control and found large and bidirectional variations in their surfaceomes (**Figure 3D**). Vimentin, a marker known to be associated with epithelial-to-mesenchymal transition (EMT) showed heightened overexpression, as well as ANPEP and fibronectin-1 (***Liu et al., 2015***). Notably, a subset of HLA molecules were downregulated in the Myc induced RWPE cells, consistent with prior findings of loss of MHC presentation in prostate cancer (***Blades et al., 1995***; ***Cornel et al., 2020***; ***Dhatchinamoorthy et al., 2021***). These findings were verified by both western blot (**Figure 3E**) and microscopy (**Figure 3F**).

**Figure 3.**
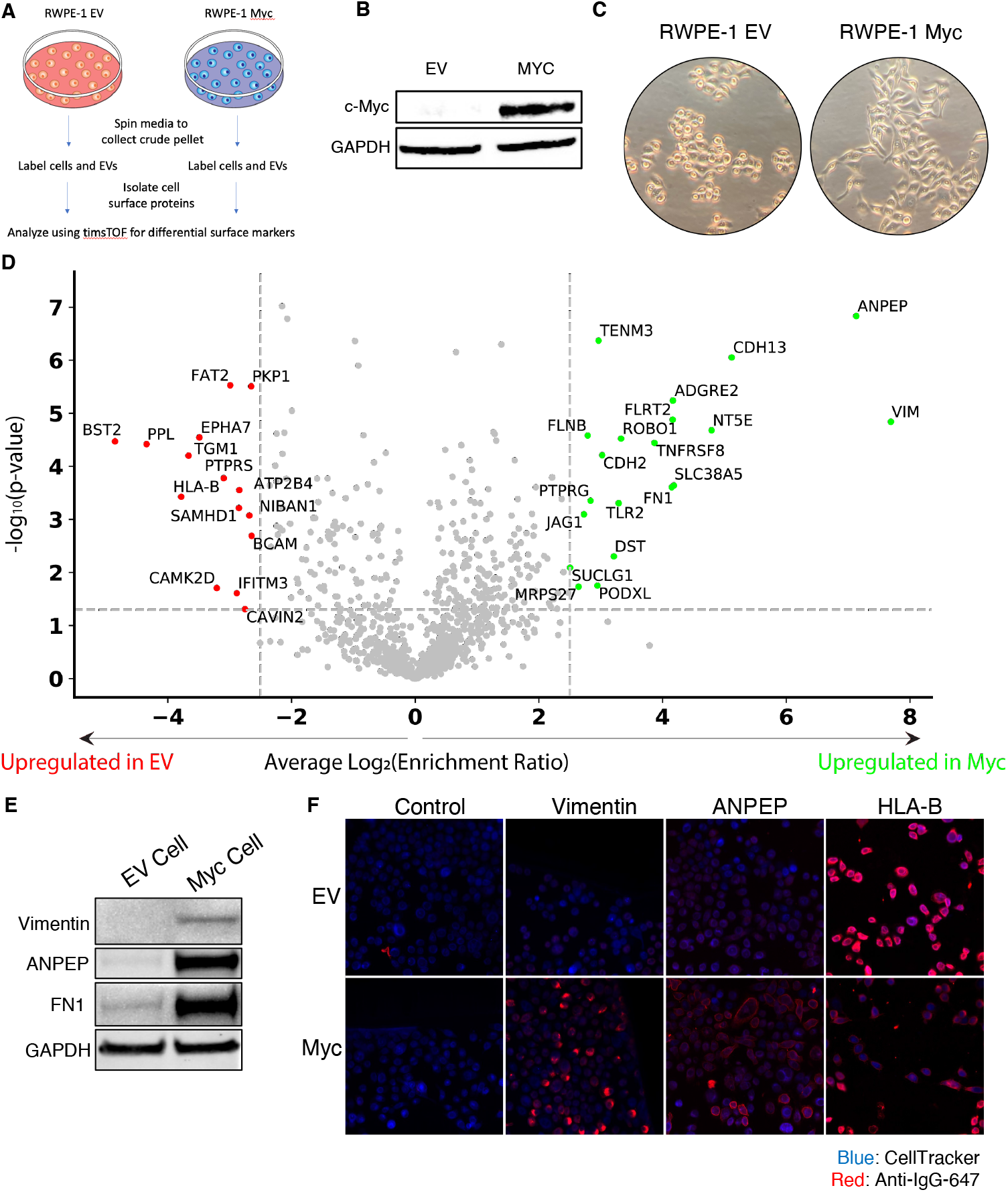
WGA-HRP identifies a number of enriched markers on Myc-driven prostate cancer cells. (A) Overall scheme for biotin labeling, and label-free quantitation (LFQ) by LC-MS/MS for RWPE-1 EV and Myc over-expression cells, and corresponding exosomes. (B) Western blot of c-Myc expression in RWPE-1 EV and Myc overexpressing cells. (C) Microscopy image depicting morphological differences between RWPE-1 EV and RWPE-1 Myc cells after 3 days in culture. (D) Volcano plot depicting the LFQ comparison of RWPE-1 EV and Myc labeled cells. Red labels indicate upregulation in the RWPE-1 EV cells over Myc cells and green labels indicate upregulation in the RWPE-1 Myc cells over EV cells. All labeled proteins are 5.6-fold enriched in either dataset between two biological replicates (p<0.05). (E) Upregulated proteins in RWPE-1 Myc cells (Vimentin, ANPEP, FN1) are confirmed by western blot. (F) Upregulated surface proteins in RWPE-1 Myc cells (Vimentin, ANPEP, FN1) are detected by immunofluorescence microscopy. The downregulated protein HLA-B by Myc over-expression was also detected by immunofluorescence microscopy.

Next, we wanted to use our WGA-HRP method to quantify cell surface proteins on exosomes derived from both normal epithelial prostate cells (RWPE-1 EV) and oncogenic Myc-induced prostate cancer cells (RWPE-1 Myc). Due to the complex process and extensive washing involved in exosome isolation, many standard labeling methods are not amenable for exosome surface labeling (**Figure 4 - Figure supplement 1**). Using WGA-HRP, we are able to biotinylate the exosomes before the sucrose gradient purification and isolation steps (**Figure 4A**). This delineated an important subset of proteins that are differentially expressed under Myc induction, which could serve as interesting targets for early-detection in patient urine or serum. This subset included fibronectin-1 (FN1), ANPEP, and ABCC1 (**Figure 4B**), which were further validated by quantitative western blotting (**Figure 4C**). A subset of these targets display similar phenotypic changes to the parent cell, suggesting that they could be biomarker candidates for non-invasive indicators of disease progression. While certain proteins are shuttled to exosomal compartments largely based off of the extent of expression in the parent cell, remarkably some proteins are singled out for exosomal packaging, indicating a pronounced differential shuttling mechanism of the proteome between cells and exosomes (**Figure 4D**). This pattern was recapitulated in the RWPE-1 EV cells and exosomes, where the majority of markers were unique to either cellular or exosomal origin (**Figure 4 - Figure supplement 2**). This is of extreme interest for not only biomarker discovery but understanding the role of exosomes in secondary disease roles, such as interfering with immune function or priming the metastatic niche (***Costa-Silva et al., 2015***).

**Figure 4.**
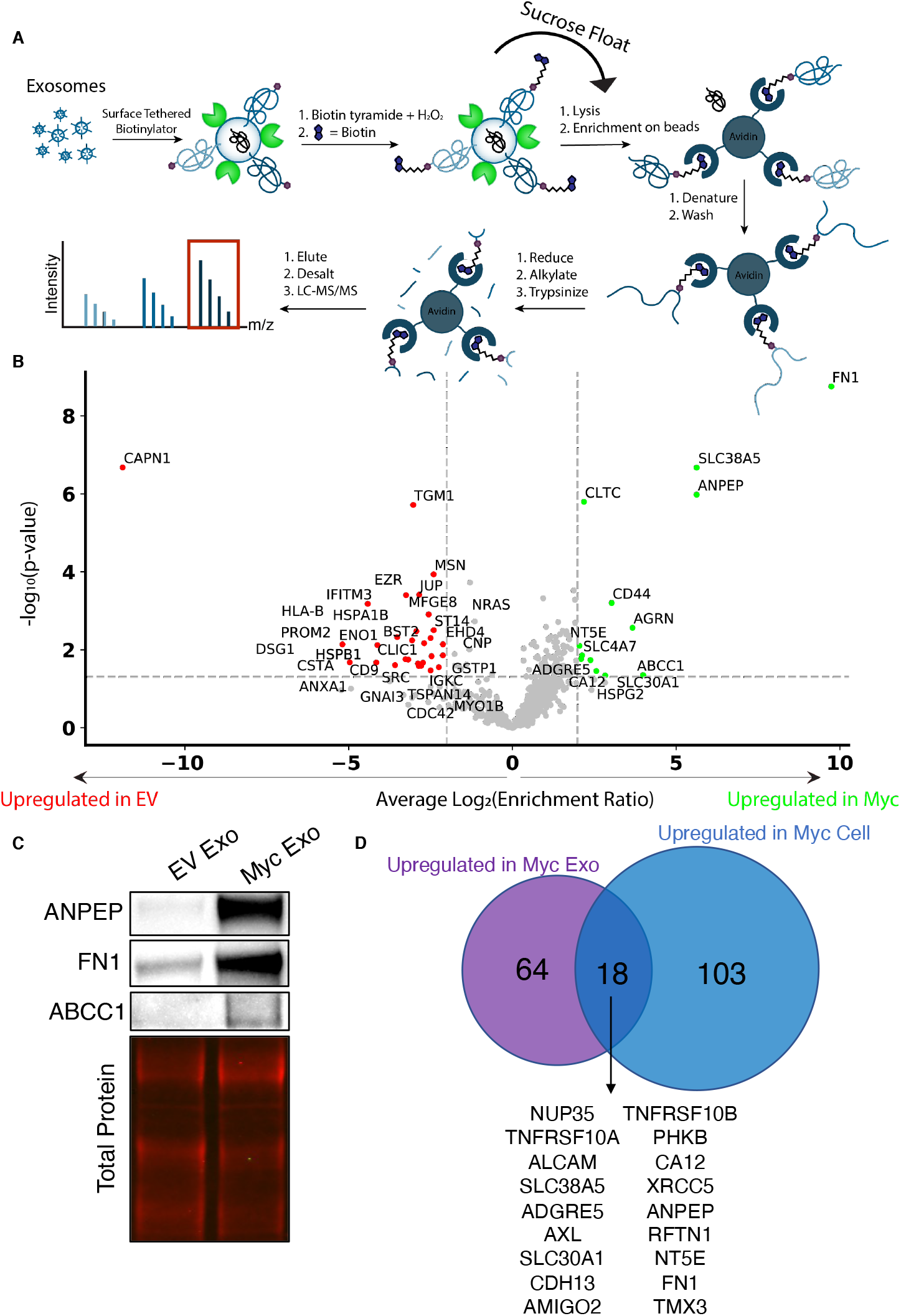
WGA-HRP identifies a number of enriched markers on Myc-driven prostate cancer exosomes. (A) Workflow of exosome labeling and preparation for mass spectrometry. (B) Volcano plot depicting label-free quantitation (LFQ) comparison of RWPE-1 Myc exosomes and EV exosomes. Proteins labeled in green are upregulated in Myc exosomes over EV exosomes and proteins labeled in red are upregulated in EV exosomes over Myc exosomes. (C) Upregulated proteins (ANPEP, FN1, ABCC1) in Myc exosomes were similarly found to be highly upregulated by western blot. (D) Venn diagram of targets upregulated on Myc-induced exosomes and Myc-induced cells compared to EV exosomes and cells, respectively.

Due to the difficulty of proteomic characterization of exosomes, our current understanding of exosomal protein shuttling remains limited. Prior proteomic exosome analysis has involved whole exosome preparations, which lacks a surface protein enrichment step (***Bandu et al., 2019***; ***Bilen et al., 2017***; ***Hosseini-Beheshti et al., 2012***). Not only is this less advantageous for the specific identification of cell surface proteins on exosomes, but it makes it impossible to compare cellular and exosome samples due to the inherent surface area-to-volume differences between cells and exosomes (***Doyle and Wang, 2019***; ***Santucci et al., 2019***). Our WGA-HRP method allows us to compare surface proteins between exosome populations, as well as between exosome and cell samples, delineating a subset of proteins that are highly upregulated in the exosomes compared to parent cell, such as ITIH4, IGSF8, and MFGE8, (**Figure 5A, 5B**) and the findings were validated by western blot (**Figure 5C**). The samples showed good overlap between replicates across all four datasets, with cellular and exosomal samples clustering by origin and oncogenic status (**Figure 4 - Figure supplement 3**). To our knowledge, this is the first experiment to wholistically characterize the surface proteome of both exosomes and parental cells. These data strongly suggest that protein triage into exosomes is a controlled process, enabling only a subset of the cell surface proteome to be shuttled to this important compartment. Our data shows that there are a variety of pan-prostate-exosome markers, notably lactadherin (MFGE8), syntenin-1 (SDCBP), serotransferrin (TF), inter-alpha-trypsin inhibitor (ITIH4), and immunoglobulin superfamily 8 (IGSF8) (**Figure 5D**), which do not seem to be Myc-specific. Indeed, when performing functional annotation clustering with the upregulated targets found in both EV and Myc exosomes, “extracellular exosome” and “extracellular vesicle” are the most significant classes given to this group of proteins (**Figure 5E**). Some of the pan-prostate exosome targets in our data have previously been linked to cancer-specific contexts, and we show here that they are also found on EV exosomes (***Shimagaki et al., 2019***; ***Tutanov et al., 2020***). Our work suggests that these markers are more broadly associated with exosomes, regardless of disease status, outlining an expanded set of targets to probe these vital compartments.

**Figure 5:**
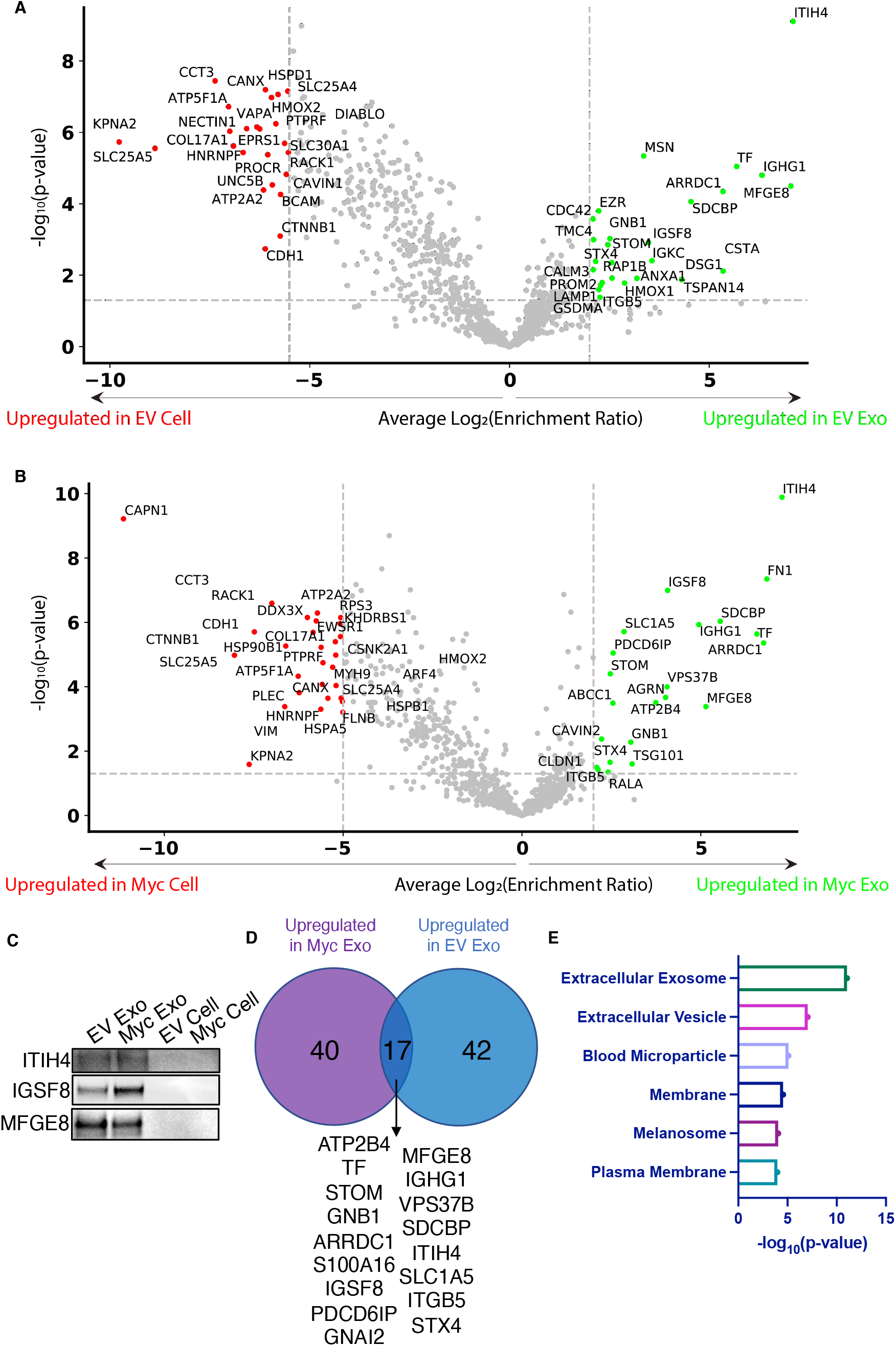
WGA-HRP identifies a number of exosome-specific markers that are present regardless of oncogene status. (A) Volcano plot depicting proteins upregulated (green) and downregulated (red) in RWPE-1 EV exosomes over EV cells. (B) Volcano plot depicting proteins upregulated (green) and downregulated (red) in RWPE-1 Myc exosomes over Myc cells. (C) Western blot showing the exosome specific marker ITIH4, IGSF8, and MFGE8. Equal amounts of total protein was loaded for each sample. (D) Overlap of 17 exosome-specific markers (>2-fold enriched). (E) Functional annotation clustering was performed using DAVID Bioinformatics Resource 6.8 to classify the 17 overlapping exosome-enriched markers.

## Discussion

The importance of understanding and characterizing cellular and exosomal membrane compartments is vital for improving our understanding of exosome biogenesis. New, improved methodologies amenable to small-scale and rapid surface proteome characterization are essential for continued development in the areas of therapeutics, diagnostics, and basic research. We sought to develop a simple, rapid surface protein labeling approach that was compatible with small sample sizes, while remaining specific to the cell surface. We took advantage of fast peroxidase enzymes and either complementary lipidated DNA technology (DNA-APEX2) or the glycan binding moiety wheat germ agglutinin (WGA-HRP) and demonstrated that tethering was much more effective than soluble addition, with increases in protein identification of between 30-90%. Additionally, we compared WGA-HRP to the existing methods, sulfo-NHS-LC-LC-biotin and biocytin hydrazide. While these alternative methods are robust, they are unable to capture time-sensitive changes, and are either plagued by low selectivity/specificity (biotin-NHS) (***Weekes et al., 2010***) or the requirement for large sample inputs (biocytin hydrazide).

There are many advantages of our new methods over the current cell surface labeling technologies. Compared to both sulfo-NHS-LC-LC-biotin and biocytin hydrazide, WGA-HRP experiments require 2 minutes instead of 30 or 120 minutes, respectively. It is also able to enrich cell surface proteins much more efficiently than sulfo-NHS-LC-LC-biotin labeling. Furthermore, NHS peptide isolation and preparation is complicated due to the reactivity of NHS chemistry towards free-amines, which blocks tryptic and LysC cleavages typically used in proteomics (***Chandler and Costello, 2016***; ***Hacker et al., 2017***).

The hydrazide method is highly effective for enriching cell surface proteins, but it is challenging for small sample sizes, due to the two-step labeling process and cell loss from the oxidation step and extensive washing. Additionally, neither biotin-NHS nor biocytin hydrazide are able to capture short time points to encompass dynamic changes at the cell surface. Due to the rapid nature of peroxidase enzymes (1-2 min), our approaches enable kinetic experiments to capture rapid changes, such as binding, internalization, and shuttling events. Another disadvantage of the hydrazide method is that it can only enrich for proteins that are glycosylated at the cell surface and it is estimated that 10-15% of cell surface proteins are not glycosylated (***Apweiler, 1999***). Glycosylation patterns also readily change during tumorigenesis, which can alter the quantification of glycan-based labeling methods, such as biocytin hydrazide (***Reily et al., 2019***). While the WGA-HRP method requires glycosylated proteins to be present to bind, it still allows for labeling of non-glycosylated proteins nearby. It is a possibility that certain cells may have low or uneven levels of glycosylation on their surfaces. In these cases, the DNA-APEX2 method can be utilized to obtain effective labeling. However, both these peroxidase-based methods require the presence of tyrosine residues (natural abundance 3.3%) to react with the biotin-tyramide radical so would not be present in all proteins (***Dyer, 1971***).

With the WGA-HRP method, we were able to compare the surfaceome of exosomes to parental cells for Myc-induced prostate cancer cells and identified proteins that were upregulated in Myc-induced cells and exosomes, as well as proteins that were differentially shuttled between exosomes and parental cells. We found a number of Myc and exosome specific markers in our study, including ANPEP, Fibronectin-1 (FN1), ABCC1, NT5E, CA12, and SLC38A5. ANPEP is a membrane-bound ectopeptidase that degrades N-termini with neutral amino acids and was found 140-fold upregulated in Myc-induced cell line compared to the EV cell line and 49-fold upregulated in the Myc-induced exosome compared to EV exosome. This peptidase has been associated with angiogenesis and cancer growth (***Guzman-Rojas et al., 2012***; ***Sørensen et al., 2013***; ***Wickström et al., 2011***). Recent studies have shown ANPEP/CD13 is systematically up-regulated on isogenic cell lines expressing proliferative oncogenes (***Leung et al., 2020***; ***Martinko et al., 2018***) or in tubular sclerosis bladder cancers (***Wei et al., 2020***), suggesting it is a commonly up-regulated in cancers. The second most differentially expressed protein between the Myc and EV samples was Fibronectin-1 (FN1), which has been shown to drive all stages of tumorigenesis (***Wang and Hielscher, 2017***). Importantly, FN1 provides an extracellular scaffold by which other matrix proteins can be deposited. Through these interactions with matrix proteins and cell-associated integrins, FN1 regulates cellular fate decisions, proliferation, and metastasis (***Efthymiou et al., 2020***).

While some proteins were present in both the exosome and cellular samples, others were only found enriched in Myc exosomes. ABCC1, also known as multi-drug resistant protein 1 (MRP1) was over 5-fold upregulated in the Myc exosomes over EV exosomes. Interestingly, this relationship was not found in the parent cells, which suggests that ABCC1 is differentially shuttled into oncogenic exosomes. The role of this protein has long been associated with imparting a chemoprotective effect on cells, due to the efflux of numerous classes of anti-cancer drugs (***Cole, 2014***).

Another such target is Agrin, which was 12-fold upregulated in the Myc exosomes over EV exosomes and has been previously seen upregulated in prostate cancer exosomes (***Hosseini-Beheshti et al., 2012***). Agrin has been shown to play an important role in the cross-talk between cancer cells and the endolethium, and contributes to ECM remodeling during tumorigenesis (***Chakraborty et al., 2020***). These targets delineate an important subset of proteins that are triaged into exosomes and could play long-range roles in promoting tumorigenesis and downstream metastasis (***Costa-Silva et al., 2015***; ***Demory Beckler et al., 2013***; ***Hoshino et al., 2015***; ***Peinado et al., 2012***).

As research shifts into analyzing native biological samples from extracellular vesicles to xenograft models or patient biopsies, it becomes increasingly important to develop sensitive, effective methods to label these small samples sizes. It is our hope that these tools will provide much needed avenues by which to pursue pressing biological questions in the areas of diagnostic and therapeutic development, as well as basic research.

## Methods and Materials

### Large-Scale APEX2 Expression, Purification, and Heme Reconstitution

APEX2 was expressed using previous methods in BL21(DE3)pLysS cells (***Howarth and Ting, 2008***). Briefly, APEX2 expression plasmid was transfected into competent BL21(DE3)pLysS cells and heat shocked for 45 seconds before being placed on ice. Cells were plated on LB/Carb plates and grown overnight at 37°C. A single colony was isolated and grown in a mixture of 30 ml of 2XYT + Carb overnight at 37°C while shaking. The overnight culture was combined with 3 L of 2XYT with Carb and placed in a 37°C shaking incubator. At an OD600 of 0.6, 100 *μ*g/ml of IPTG was added and the temperature of the incubator was lowered to 30°C. Cells were allowed to incubate for 3.5 hours and spun down at 6,000xg for 20 minutes. Cell pellet was resuspended in protease inhibitor containing resuspension buffer (5 mM Imidazole, 300 mM NaCl, 20 mM Tris pH=8) and mixed thoroughly. The mixture was sonicated at 50% for 5 seconds on:15 seconds off for 5 minutes on ice to avoid bubble formation. Lysate was mixed by inversion at 4°C for 15 minutes and spun down at 19,000xg for 20 minutes. The slurry was introduced to 5 ml of washed Nickel resin slurry and allowed to bind by gravity filtration. The beads were washed 3x with wash buffer (30 mM Imidazole, 300 mM NaCl, 20 mM Tris pH=8) and eluted in 5 ml of elution buffer (250 mM Imidazole, 300 mM NaCl, 20 mM Tris pH=8) before undergoing buffer exchange into PBS.

Enzyme underwent heme reconstitution as per previous methods (***Cheek et al., 1999***). Briefly, 50 mg of hemin-Cl (Sigma) was diluted in 2.0 mL of 10 mM NaOH. The mixture was thoroughly resuspended, then diluted further using 8.0 mL of 20 mM KPO4, pH 7.0, and vortexed extensively. Mixture was spun down at 4,000xg 2x to get rid of insoluble hemin. APEX2 was diluted 1:2 in 20 mM KPO4. 6 ml of heme stock was added to 2 ml of APEX over 20 minutes and allowed to rotate at 4°C wrapped in tin foil for 3 hours. The mixture was introduced to a column with 20 ml of DEAE Sepharose pre-equilibrated in 20 mM KPO4, pH 7.0 buffer. Enzyme was eluted using 100 mM KPO4 and spin concentrated. To verify complete reconstitution, absorbance was measured at 403 and 280 nm. A403/280 > 2.0 is considered sufficient for reconstitution. The isolated protein was flash frozen and stored at −80°C for long-term storage. Each batch of enzyme was run out on a 4-12% Bis-Tris gel to confirm purity (**Figure 1-Figure supplement 1**).

### APEX2 DNA labeling protocol

APEX2 was incubated at 50 μM with 40 molar equivalents of maleimide-DBCO for 5 hours at room temperature in PBS. The reaction was desalted with Zeba columns (7 kDa cutoff). 2.5 molar equivalents of Azido-DNA was added to the reaction and incubated at 4°C overnight. Successful conjugation was monitored by LC-MS before the mixture was purified by nickel column.

### Cell culture

Expi293 suspension cells were maintained in Expi293 media (Thermo, A1435101) and rotating at 125 rpm in a 37°C incubator with 8% CO2.Cells were split every 3 days by diluting into new media. Adherent PaTu8902 and KP-4 cells were grown in pre-warmed Iscove’s Modified Dulbecco’s Media (IMDM) supplemented with 10% FBS (Gemini Bio-Products, 100-106) and 5% Penicillin/Streptomycin (Thermo Fisher Scientific, 15-140-122) at 37°C in a 5% CO2-humidified incubator. Adherent RWPE-1 prostate cells were grown in complete keratinocyte-SFM (Thermo; 17005-042) supplemented with bovine pituitary extract (BPE), recombinant EGF, and 5% penicillin/streptomycin at 37°C in a 5% CO2-humidified incubator. The media was exchanged every two days. For splitting, cells were lifted with 0.05% Trypsin (Life Technologies) and quenched with 5% FBS before spinning down cells to remove residual trypsin and FBS. Cells were then plated in pre-warmed complete keratinocyte-SFM media.

### Microscopy

Cells were plated at a density of 15,000 cells per well in a 96-well clear bottom plate (Greiner Bio-One, 655090) pre-treated with poly-D-lysine (Thermo Scientific, A3890401). Cells were allowed 48 hours to reattach and grow undisturbed. Cells were washed 3x in cold PBS. For DNA-APEX2, 100 μl of 0.5 μM enzyme solution was combined with anchor and co-anchor at a final concentration of 1 μm. For all other enzymes, enzyme was combined with PBS at a final concentration of 0.5 μM. For sugar blocking studies, 100 μl of dilution enzyme solution (0.5 μM) was combined with 100 mg/ml N-acetyl-D-glucosamine (Sigma Aldrich, A3286-5G). Cells were allowed to sit on ice for 5 minutes to allow WGA to bind fully, as labeling was not altered by increased incubation time (**Figure 1 - Supplementary Figure 3**). Biotin tyramide (Sigma Aldrich, SML2135-50MG) was added to cells with a final concentration of 500 μM before adding 1 mM of H_2_O_2_. Reaction was allowed to continue for 2 minutes before rinsing cells 3x with 1X quench buffer (10 mM sodium ascorbate + 5 mM Trolox + 1 mM sodium pyruvate). The cells were rinsed 2x with PBS and crosslinked with 4% PFA for 10 minutes at RT. Cells were washed 3x with PBS before introduction to 1:100 primary antibody. Primary antibodies used were HisTag-650 (Invitrogen, MA1-21315-D650), Streptavidin-488 (Thermo Fisher Scientific, S-11223), biotin-conjugated anti-HRP (Rockland, 200-4638-0100), ANPEP (R&D Systems, AF3815), vimentin (Cell Signaling Technology, 5741S), FN1 (Abcam, ab2413), and HLA-B (Protein-Tech, 17260-1-AP). Cells were washed 3x in PBS and imaged on an IN Cell Analyzer 6500. Images were processed in Fiji using the Bio Formats plugin (***Linkert et al., 2010***; ***Schindelin et al., 2012***).

### Cell-tethered APEX2, soluble APEX2, cell-tethered WGA-HRP and soluble HRP cell surface labeling

Cultured cells were grown for 3 days in tissue culture plates and dissociated by addition of versene (PBS + 0.05% EDTA). Cells were washed 3x in PBS (pH 6.5), resuspended in PBS (pH 6.5) and aliquoted to 500,000 cells per sample. Samples were resuspended in 100 μL of PBS (pH 6.5). For anchored APEX2 samples, lipidated anchor DNA was allowed to bind for 5 minutes at 1 μM on ice, followed by 1 μM of lipidated co-anchor DNA on ice for 5 minutes. 0.5 μM DNA-labeled APEX2 was allowed to bind on cells for 5 minutes before final wash with PBS (pH 6.5). For soluble APEX2, WGA-HRP, and soluble HRP samples, cells were resuspended in 0.5 μM of the corresponding enzyme. WGA-HRP was allowed to bind to cells for 5 minutes on ice. Biotin tyramide was added at a final concentration of 500 μM and mixed thoroughly, before the addition of 1 mM H_2_O_2_. Cells underwent labeling in a 37°C incubator for 2 minutes before being quenched with 5 mM Trolox/10 mM Sodium Ascorbate/1 mM Sodium Pyruvate. Cells were washed 2x in quench buffer and spun down. The pellet was either further processed for flow cytometry, western blot, or flash frozen in liquid nitrogen for mass spectrometry.

### On plate WGA-HRP cell surface labeling

KP-4 cells were grown on a 6 cm tissue culture treated plate and washed 3x with PBS (pH 6.5). 2 mL of 0.5 μM WGA-HRP in PBS (pH 6.5) was added to the plate, followed by biotin tyramide (0.5mM final concentration) and H_2_O_2_ (1mM final concentration). After a 2 minute incubation at 37°C, the cells were washed 2x with 5 mM Trolox/10 mM Sodium Ascorbate/1 mM Sodium Pyruvate quenching solution. The cells were washed 1x with PBS before being lifted with versene (PBS + 0.05% EDTA). Once lifted, the cells were washed once with PBS and subsequentially processed for flow cytometry analysis.

### Biocytin hydrazide cell surface labeling

Cultured cells were grown for 3 days in tissue culture plates and dissociated by addition of versine (PBS + 0.05% EDTA). Cells were washed 3x in PBS (pH 6.5), resuspended in PBS (pH 6.5) and aliquoted to 1.5 million cells per sample. Samples were resuspended in 100 μL of PBS (pH 6.5) and fresh sodium periodate (1 μL of a 160 mM solution) was added to each sample. The samples were mixed, covered in foil, and incubated rotating at 4°C for 20 minutes. Following three washes with PBS (pH 6.5), the samples were resuspended in 100 μL of PBS (pH 6.5) with the addition of 1 μL of aniline (diluted 1:10 in water) and 1 μL of 100 mM biocytin hydrazide (Biotium, 90060). The reaction proceeded while rotating at 4°C for 90 minutes. The samples were then washed 2x with PBS (pH 6.5) and spun down. The pellet was either further processed for flow cytometry, western blot, or flash frozen in liquid nitrogen for mass spectrometry.

### Sulfo-NHS-LC-LC-biotin cell surface labeling

Cultured cells were grown for 3 days in tissue culture plates and dissociated by addition of versine (PBS + 0.05% EDTA). Cells were washed 3x in PBS (pH 7.4), resuspended in PBS (pH 8) and aliquoted to 1.5 million cells per sample. Samples were resuspended in 50 μL of PBS (pH 8). An aliquot of EZ-Link Sulfo-NHS-LC-LC-Biotin (ThermoFisher, 21338) was resuspended in 150 μL of PBS (pH 8). 7.5 μL was added to each cell sample and the reaction proceeded rotating at 4°C for 30 minutes. The reaction was quenched by the addition of 2.5 μL of 1M Tris (pH 8.0). The samples were washed 2x in PBS (pH 7.4) and spun down. The pellet was either further processed for flow cytometry, western blot, or flash frozen in liquid nitrogen for mass spectrometry.

### Flow cytometry for cell surface biotinylation

After labeling and quench washes, cells were washed once with PBS + 2% BSA to inhibit nonspecific binding. Samples were then incubated with 100 μL Streptavidin-Alexa Fluor 647 (Thermo Fischer, 1:100 in PBS + 2% BSA). Following a 30-minute incubation at 4°C while rocking, samples were washed three times with PBS + 2% BSA. Samples were analyzed in the APC channel and quantified using a FACSCanto II (BD Biosciences). All flow cytometry data analysis was performed using FlowJo software.

### RWPE-1 exosome isolation and labeling protocol

Exosomes were isolated as previously described (***Poggio et al., 2019***). Briefly, the day prior to exosome isolation, media was replaced with BPE-free keratinocyte-SFM media. For vesicle enrichment, media was isolated after two days in BPE-free media and centrifuged at 300 x g for 10 minutes at RT, followed by 2,000 x g for 20 minutes at 4°C. Large debris was cleared by a 12,000 x g spin for 40 minutes at 4°C. The pre-cleared supernatant was spun a final time at 100,000 x g at 4°C for 1 hr to pellet extracellular vesicles. Isolated extracellular vesicles were brought up in 50 μl of PBS with 0.5 μM of WGA-HRP and mixture was allowed to bind on ice for 5 minutes. WGA-HRP bound vesicles were placed on a shaker (500 rpm) at 37 °C before the addition of biotin tyramide (0.5 mM final concentration) and H_2_O_2_ (1 mM final concentration). Vesicles underwent labeling for 2 minutes before being quenched with 5 mM Trolox/10 mM Sodium Ascorbate/1 mM Sodium Pyruvate. Biotinylated exosomes were purified from extracellular vesicles by further centrifugation on a sucrose gradient (20-60%) for 16 hours at 4°C at 100,000xg.

### Western blot protocol

Cultured cells were grown in 15 cm^2^ tissue culture plates and dissociated by addition of versine (PBS + 0.05% EDTA). Cells were washed in PBS (pH 6.5) and resuspended in 100 μl PBS (pH 6.5) at a concentration of 10 million cells/ml in PBS (pH 6.5). Cells were labeled, reaction was quenched with 1X NuPage Loading Buffer, and immediately boiled for 5 minutes. To enable proper addition of lysate to gel wells, the mixture was thinned with addition of nuclease, and the disulfides were reduced with BME. The samples were subjected to electrophoresis in a 4-12% NuPage Gel until the dye front reached the bottom of the gel cast. For cell and exosome blots, equal amounts of sample was prepared in 1X NuPage Loading Buffer with BME and boiled for 5 minutes. Samples were loaded and subjected to electrophoresis in a 4-12% NuPage Gel until the dye front reached the bottom of the gel cast. Prepared gels were placed in iBlot2 transfer stacks and transferred using the P0 setting on the iBlot 2 Gel Transfer Device. The PVDF membrane was blocked in TBS Odyssey Blocking buffer for 1 hour at RT. Membranes were washed in TBST and incubated with Strepavidin-800 (1:10,000 dilution, Licor, 926-32230) for 30 minutes or in TBS Odyssey Blocking buffer + 0.1% Tween 20. Membranes were washed in TBST 3x with a final wash in water. Membranes were visualized using an Odyssey DLx imager.

For cell and exosome blots, equal amounts of sample was prepared in 1X NuPage Loading Buffer with BME and boiled for 5 minutes. Samples were loaded and subjected to electrophoresis in a 4-12% NuPage Gel until the dye front reached the bottom of the gel cast. Prepared gels were placed in iBlot2 transfer stacks and transferred using the P0 setting on the iBlot 2 Gel Transfer Device. The PVDF membrane was blocked in TBS Odyssey Blocking buffer for 1 hour at RT. Membranes were washed in TBST and incubated overnight in primary antibody at 4°C in TBS Odyssey Blocking buffer + 0.1% Tween 20 while shaking. Primary antibodies used were ANPEP (R&D Systems, AF3815), FN1 (Abcam, ab2413), ABCC1 (Cell Signaling Technology, 72202S), ITIH4 (Atlas Antibodies, HPA003948), MFGE8 (Thermo Scientific, PA5-82036), IGSF8 (R&D Systems, AF3117-SP). Mem-branes were washed in 3x TBST before introduction to a 1:10,000 dilution of secondary antibody in TBS Odyssey Blocking buffer + 0.1% Tween 20 for 1 hour at room temperature while shaking. Secondary antibodies used were Goat Anti-Rabbit HRP (Thermo Scientific, 31460) and Rabbit Anti-Sheep HRP (Thermo Scientific, 31480). Blots were imaged after 5 minutes in the presence of Super-Signal West Pico PLUS Chemiluminescent Substrate (Thermo Fisher Scientific, 34577) and imaged using a ChemiDoc XRS+.

### Proteomic sample preparation

Frozen cell and exosome pellets were lysed using 2X RIPA buffer (VWR) with protease inhibitor cocktail (Sigma-Aldrich; St. Louis, MO) at 4°C for 30 mins. Cell lysate was then sonicated, clarified, and incubated with 100 μl of neutravidin agarose slurry (Thermo Fisher Scientific) at 4°C for 1 hr. The bound neutravidin beads were washed in 2 ml Bio-spin column (Bio-Rad, 732-6008) with 5 ml RIPA buffer, 5 ml high salt buffer (1M NaCl, PBS pH 7.5), and 5 ml urea buffer (2M urea, 50mM ammonium bicarbonate) to remove non-specific proteins. Beads were allowed to fully drain before transferring to a Low-bind Eppendorf Tube (022431081) with 2M Urea. Sample was spun down at 1,000xg and aspirated to remove excess liquid. Samples were brought up in 100 μl of 4M Urea digestion buffer (50 mM Tris pH 8.5, 10 mM TCEP, 20 mM IAA, 4 M Urea) and 2 μg of total reconstituted Trypsin/LysC was added to the sample before incubating for 2 hours at RT. To activate the trypsin, mixture was diluted with 200 μl of 50 mM Tris pH 8.5 to a final Urea concentration of below 1.5 M. The mixture was covered and allowed to incubate overnight at RT. The mixture was isolated from the beads by centrifugation (Pierce; 69725) before being acidified with 10% TFA until pH of 2 was reached. During this time, a Pierce C18 spin column was prepared as per manufacturing instructions. Briefly, C18 resin was washed twice with 200 μl of 50% LC-MS/MS grade ACN. The column was equilibrated with two 200μl washes of 5% ACN/0.5% TFA. The pre-acidified sample was loaded into the C18 column and allowed to fully elute before washing twice with 200μl washes of 5% ACN/0.5% TFA. One final wash of 200 μl 5% ACN/1% FA was done to remove any residual TFA from the elution. Samples were eluted in 70% ACN, dried, and dissolved in 0.1% formic acid, 2% acetonitrile prior to LC-MS/MS analysis. Peptides were quantified using Pierce Quantitative Colorimetric Peptide Assay (Thermo Fisher Scientific, 23275).

### LC-MS/MS

Liquid chromatography and mass spectrometry was performed as previously described (***Meier et al., 2020***). Briefly, approximately 200 ng of peptides were separate using a nanoElute UHPLC system (Bruker) with a pre-packed 0.75mm x 150mm Acclaimed Pepmap C18 reversed phase column (120 A pore size, IonOpticks) and analyzed on a timsTOF Pro (Bruker) mass spectrometer. Peptides were separated using a linear gradient of 2-34% solvent B (Solvent A: 0.1% formic acid, solvent B: 80% acetonitrile, 0.1% formic acid) over 100 mins at 400 nL/min. Data-dependent acquisition was performed with parallel accumulation-serial fragmentation (PASEF) and trapped ion mobility spectrometry (TIMS) enabled with 10 PASEF scans per topN acquisition cycle. The TIMS analyzer was operated at a fixed duty cycle close to 100% using equal accumulation and ramp times of 100 ms each. Singly charged precursors were excluded by their position in the m/z–ion mobility plane, and precursors that reached a target value of 20,000 arbitrary units were dynamically excluded for 0.4 min. The quadrupole isolation width was set to 2 m/z for m/z < 700 and to 3 m/z for m/z > 700 and a mass scan range of 100-1700 m/z. TIMS elution voltages were calibrated linearly to obtain the reduced ion mobility coefficients (1/K0) using three Agilent ESI-L Tuning Mix ions (m/z 622, 922 and 1,222).

### Data Processing

Briefly, for general database searching, peptides for each individual dataset were searched using PEAKS Online X version 1.5 against the plasma membrane annotated human proteome (Swiss-prot GOCC database, August 3, 2017 release). We acknowledge the identification of a number of proteins not traditionally annotated to the plasma membrane, which were published in the final Swissprot database used. Enzyme specificity was set to trypsin + LysC with up to two missed cleavages. Cysteine carbamidomethylation was set as the only fixed modification; acetylation (N-term) and methionine oxidation were set as variable modifications. The precursor mass error tolerance was set to 20 PPM and the fragment mass error tolerance was set to 0.03 Da. Data was filtered at 1% for both protein and peptide FDR. For comparative label-free quantification of cellular and exosomal samples, datasets were searched using MaxQuant and further analysis was performed using Perseus. Enzyme specificity was set to trypsin + LysC with up to two missed cleavages. Cysteine carbamidomethylation was set as the only fixed modification; acetylation (N-term) and methionine oxidation were set as variable modifications. The precursor mass error tolerance was set to 20 PPM and the isotope mass error tolerance was set to 0.005 Da. Data was filtered at 1% for both protein and PSM FDR. For further analysis in Perseus, proteins were removed with less than 2 unique peptides. Contaminants were removed. All peak areas were log2(x) transformed and missing values were imputed separately for each sample using the standard settings (width of 0.3, downshift of 1.8). Significance was based off of a standard unpaired Student t test with unequal variances across all four replicates. Reported peak area values represent the averages of all four replicates. The mass spectrometry proteomics data have been deposited to the ProteomeXchange Consortium via the PRIDE (***Perez-Riverol et al., 2019***) partner repository with the dataset identifier PXD028523.

## Acknowledgments

We acknowledge the members of the Wells lab for their support. Special thanks to Alice Ting, PhD and Tess Branon, PhD for their helpful advice regarding enzyme purification. We would also like to acknowledge and thank the lab of Zev Gartner, PhD for providing materials and methods for DNA-lipid tethering. J.A.W. was supported by generous funding from the Chan Zuckerberg Biohub Investigator Program, the Harry and Dianna Hind Professorship, NIH (R35GM122451), and NCI (R01CA248323). The NIH F31 Ruth L Kirschstein National Research Service Award (1F31CA247527) supported L.L.K. The National Science Foundation Graduate Research Fellowship Program (1650113) supported S.K.E. R.B. and J.Y. were supported by funding from the NIH (U01CA244452).

---

**Figure 1–Figure supplement 1.**
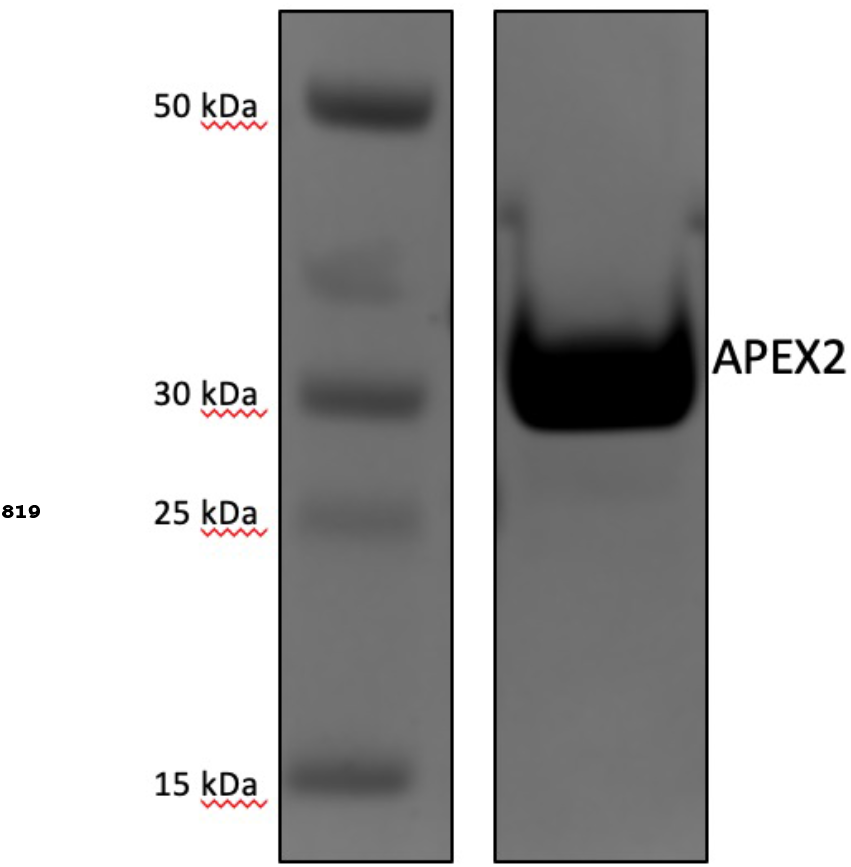
Expression, purification, and validation of APEX2 enzyme. His-tagged APEX2 was expressed in BL21(DE3)pLysS cells and purified by a nickel column. 10 μg of purified enzyme was run out on a 4-12% Bis-Tris gel to confirm purity.

**Figure 1–Figure supplement 2.**
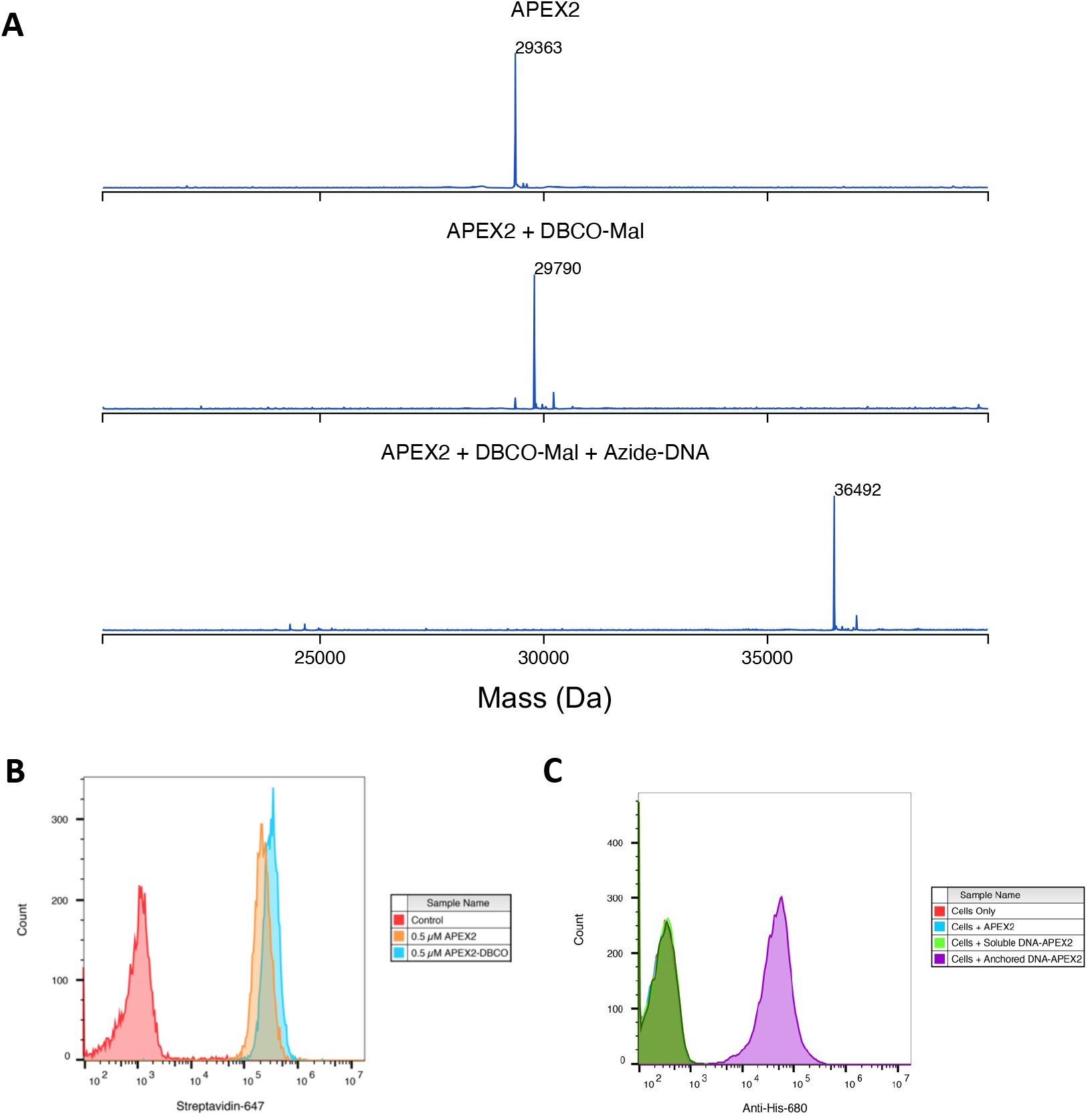
Labeling and efficacy of APEX2 with DNA. (A) APEX2 was first conjugated with DBCO-Maleimide (DBCO-Mal) reagent at 40 equivalents for 5 hours (80% conversion to the singly labeled product). Following desalting, 3 equivalences of Azide-DNA was added to the conjugate and purified by a Ni2+ column. Both reactions were monitored by LC-MS as shown. (B) 500,000 Expi293 cells were labeled with 0.5 μM purified APEX2 and DBCO-labeled APEX2 for 2 min. Extent of biotinylation of target cells was quantified by flow cytometry staining with streptavidin-647. (C) The DNA-APEX2 conjugate was shown to be tethered in the presence of the lipidated DNA (purple) and not in the absence (green), as detected by an Anti-His 680 antibody. Unlabeled APEX2 (blue) additionally did not result in a signal shift

**Figure 1–Figure supplement 3.**
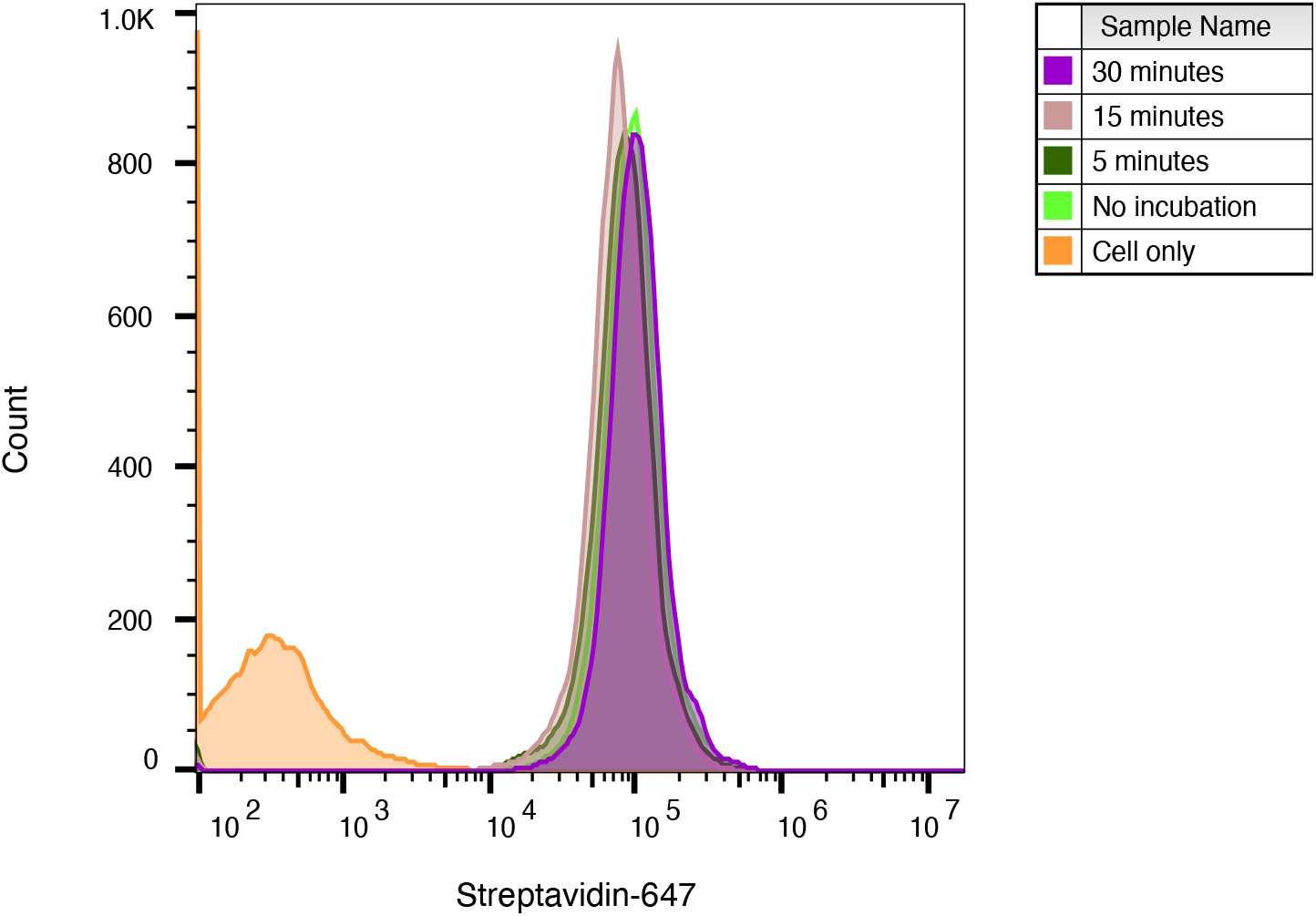
WGA-HRP pre-incubation time on cells has no effect on labeling efficiency. WGA-HRP was incubated on Expi293 cells for 0-30 min to determine optimal incubation time on ice before labeling. All tested times resulted in similar cell surface biotinylation efficiencies and signified that no incubation time was needed.

**Figure 2–Figure supplement 1.**
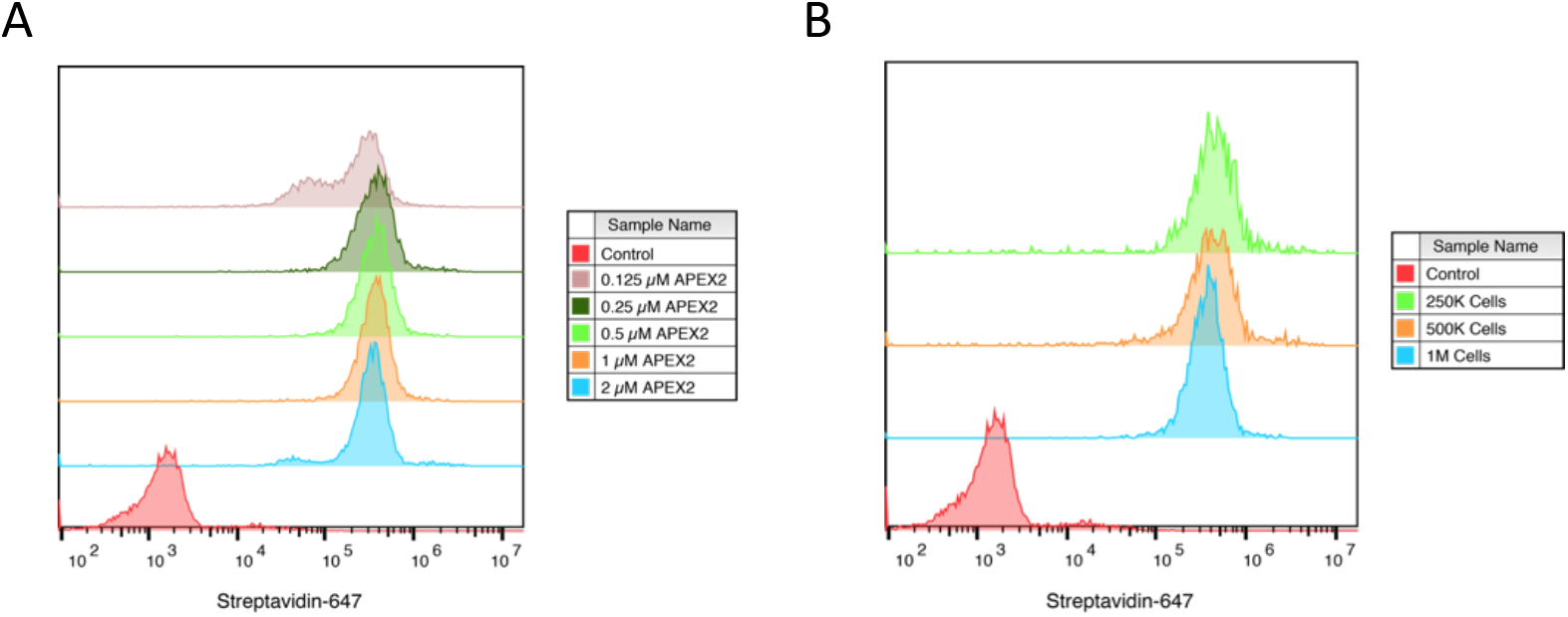
Optimization of APEX2 concentrations on cell by flow cytometry. (A) 500,000 Expi293 cells were labeled for 2 min with increasing amounts of purified APEX2 enzyme and extent of labeling was quantified by flow cytometry staining with streptavidin-647. (B) Varying numbers of Expi293 cells were labeled for 2 min with 0.5 μM APEX2 to test range of cell numbers for labeling.

**Figure 2–Figure supplement 2.**
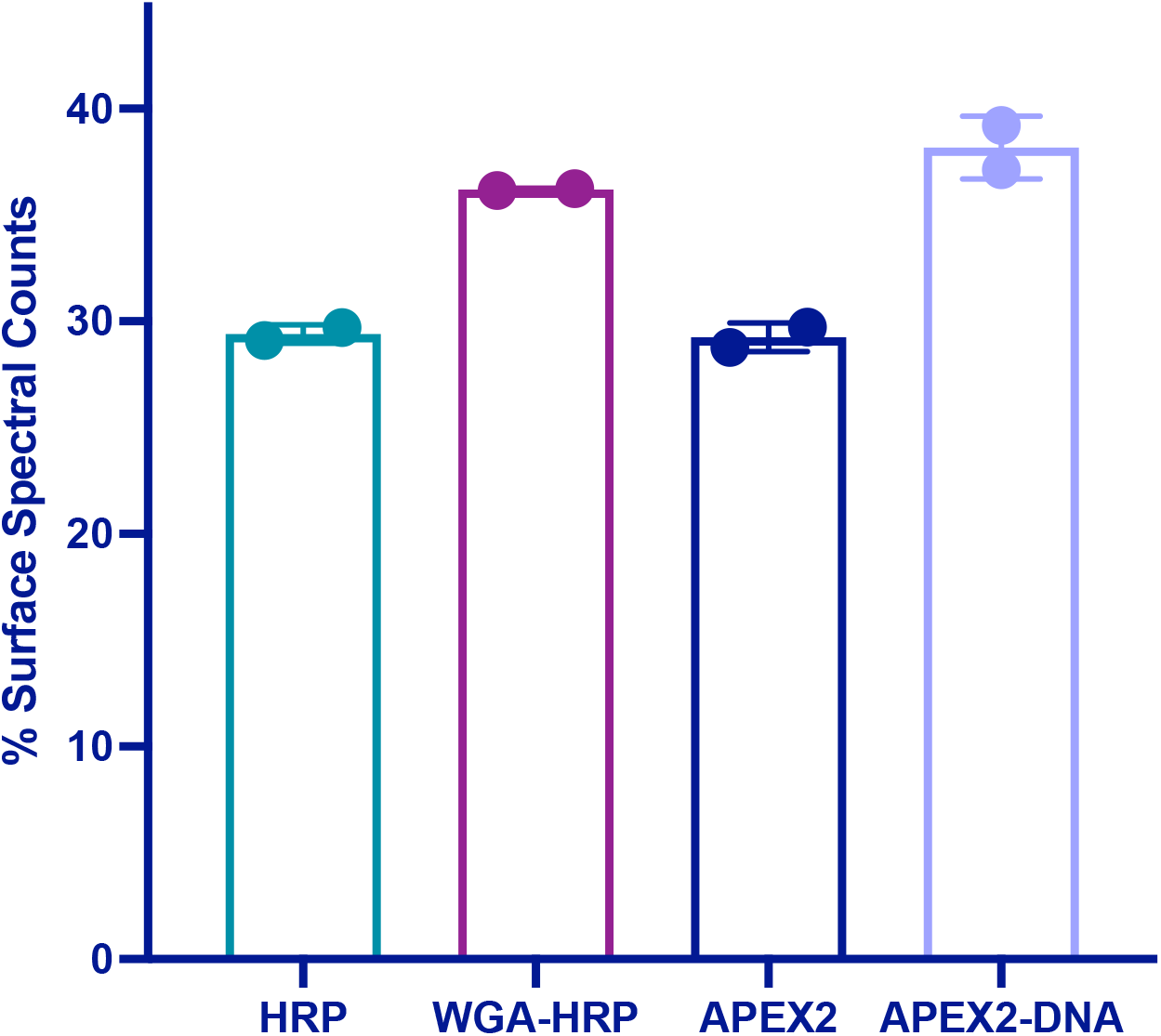
Percentage of spectral counts from plasma membrane-derived peptides across non-tethered and tethered cellular labeling experiments. The percentage of total spectral counts detected from surface-derived peptides were divided by total spectral counts detected across the entire human proteome to return a surface peptide percentage score.

**Figure 2–Figure supplement 3.**
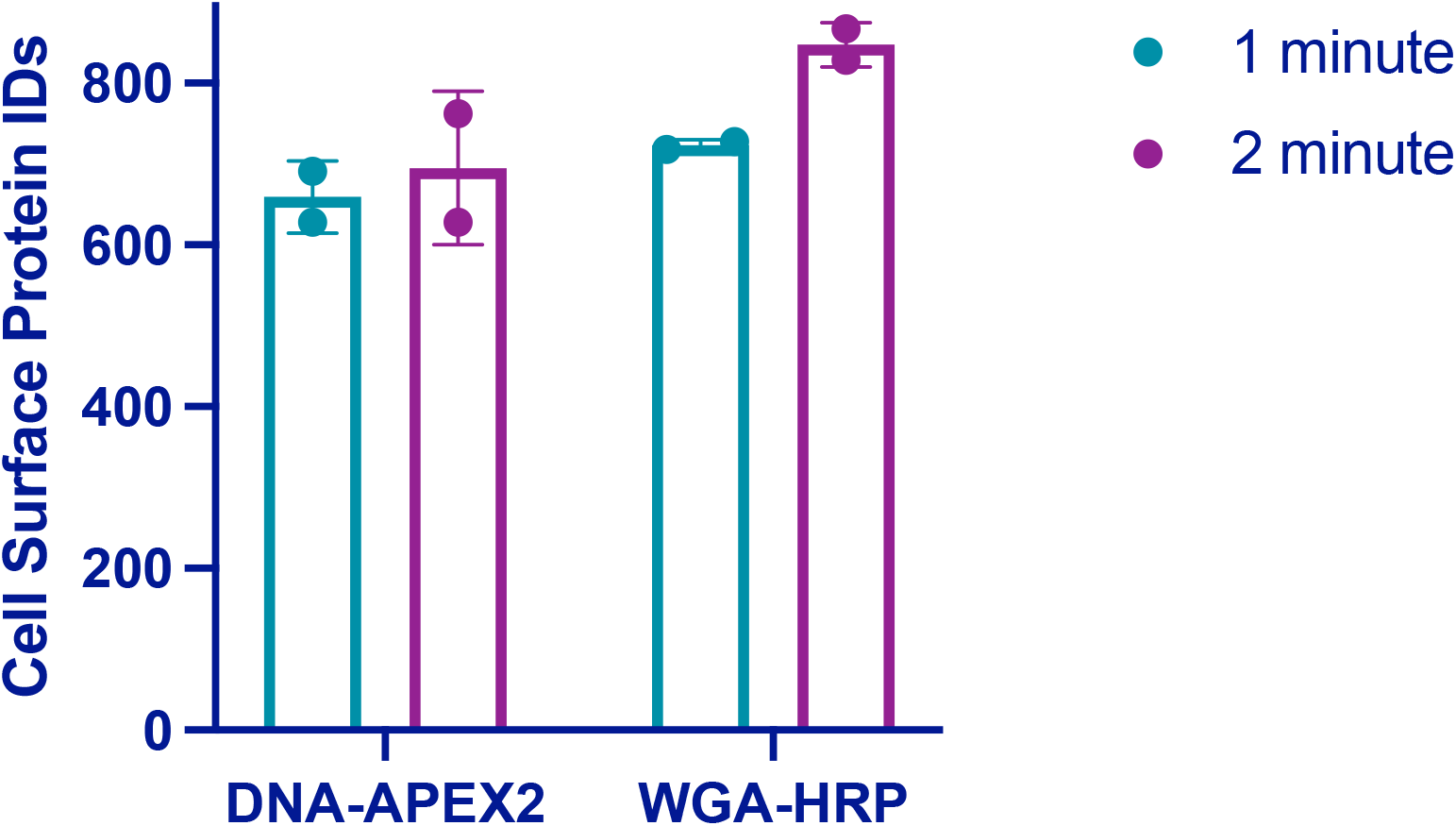
Total plasma membrane protein identifications for DNAAPEX2 and WGA-HRP labeling experiments as function of time. 500,000 PaTu8902 pancreatic cancer cells were labeled with either 0.5 μM DNA-APEX2 or 0.5 μM WGA-HRP for 1 or 2 minutes at 37°C. After cell surface enrichment and mass spectrometry analysis, the plasma membrane derived protein identification were totaled.

**Figure 2–Figure supplement 4.**
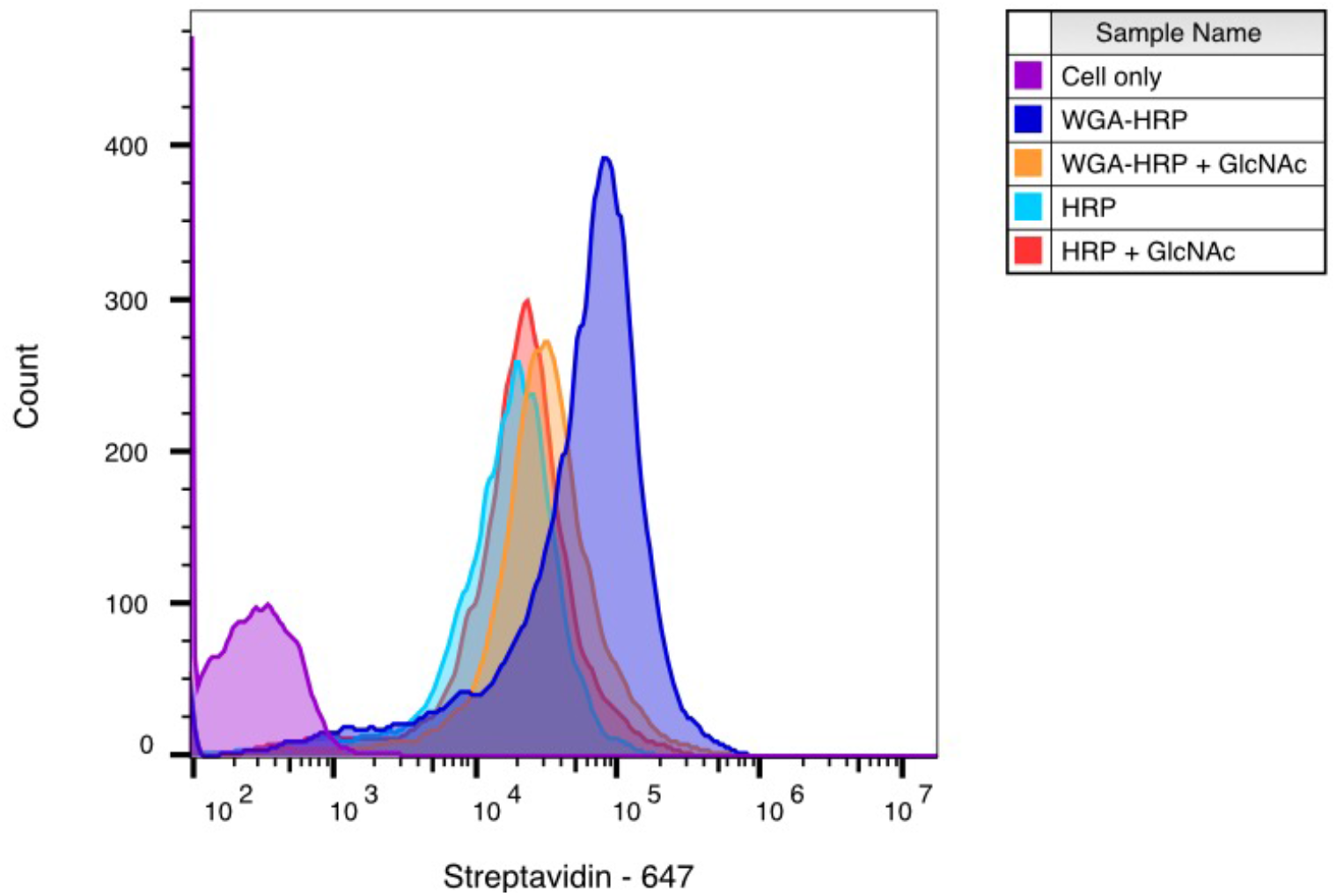
WGA-HRP labeling is N-acetcylglucosamine (GlcNAc) dependent. Biotinylation of RWPE-1 Myc cells with WGA-HRP was determined with (orange) and without (dark blue) 100 mg/mL GlcNAc. There is a significant leftward shift in the degree of labeling in the absence of competing GlcNAc, demonstracting that the enhanced labelling by WGA-HRP is GlcNAc dependent. The degree of labelling is similar to soluble HRP, as shown in light blue. Importantly, presence of GlcNAc in solution did not generally affect HRP labeling as seen by the control in red.

**Figure 2–Figure supplement 5.**
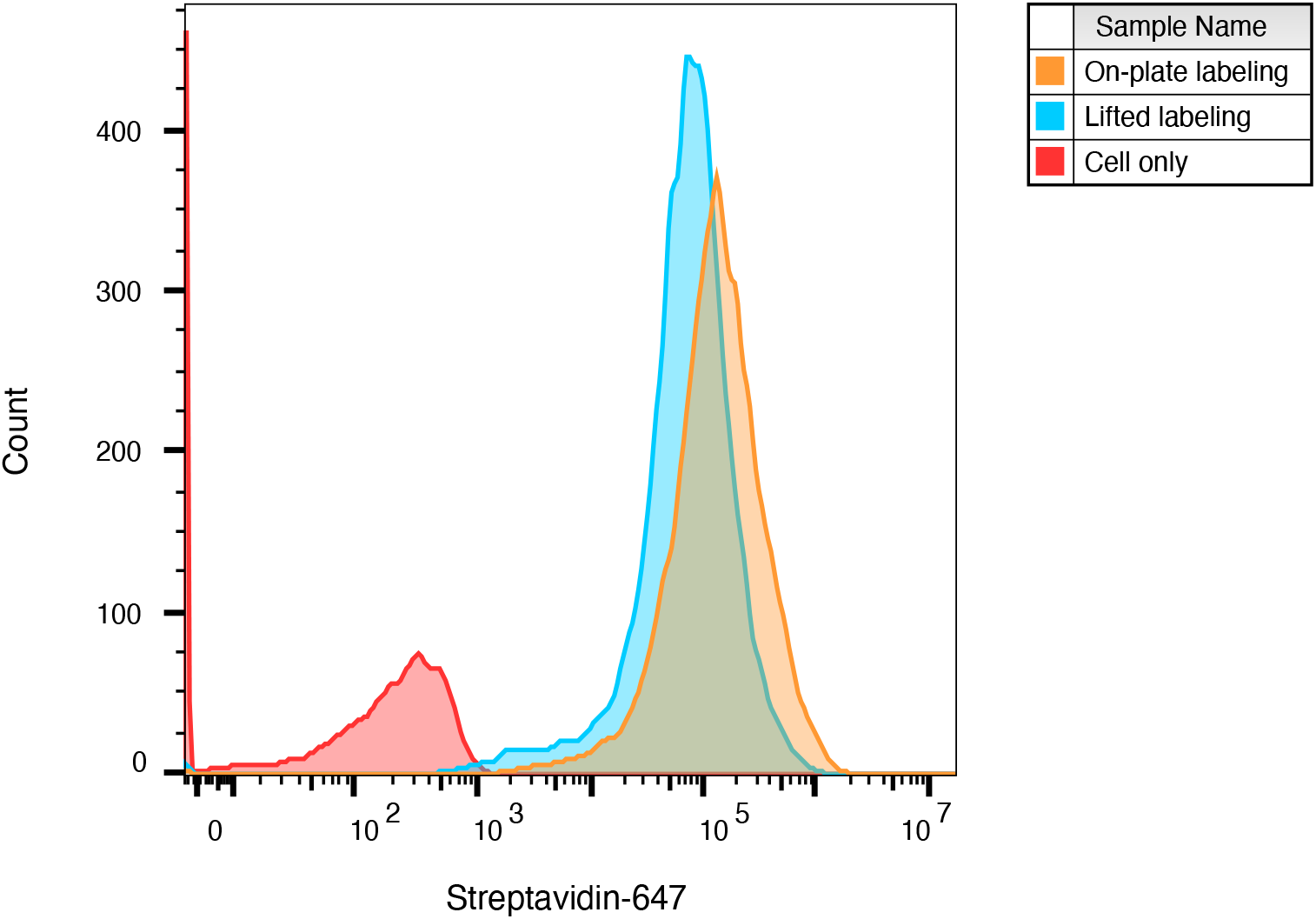
WGA-HRP can be used to label adherent cells on-plate. Cell surface labeling was compared between labeling adherent cells on a tissue culture plate vs. lifting cells and then performing labeling. Cell surface biotinylation was detected by streptavidin-Alexa Fluor 647.

**Figure 3–Figure supplement 1.**
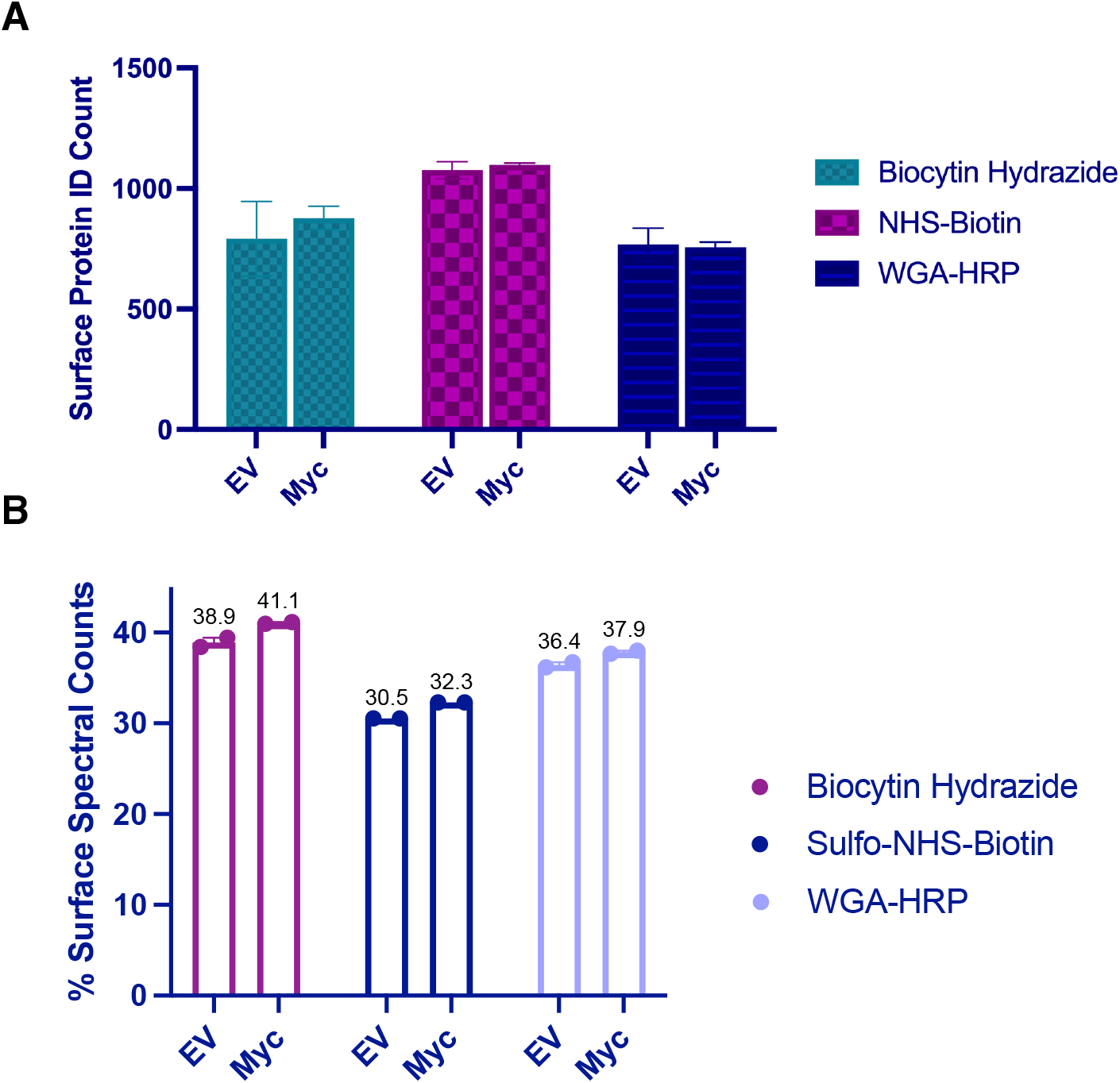
Comparison of replicates for different mass spectrometry methods. (A) The top three methods (Biotin-NHS, Biocytin Hydrazide, and WGA-HRP) were compared for their ability to identify cell surface proteins on 1.5 M RWPE-1 EV and RWPE-1 Myc cells by LC-MS/MS. (B) The percentage of total spectral counts detected from surface peptides were divided by total spectral counts detected to return a surface peptide percentage score.

**Figure 3–Figure supplement 2.**
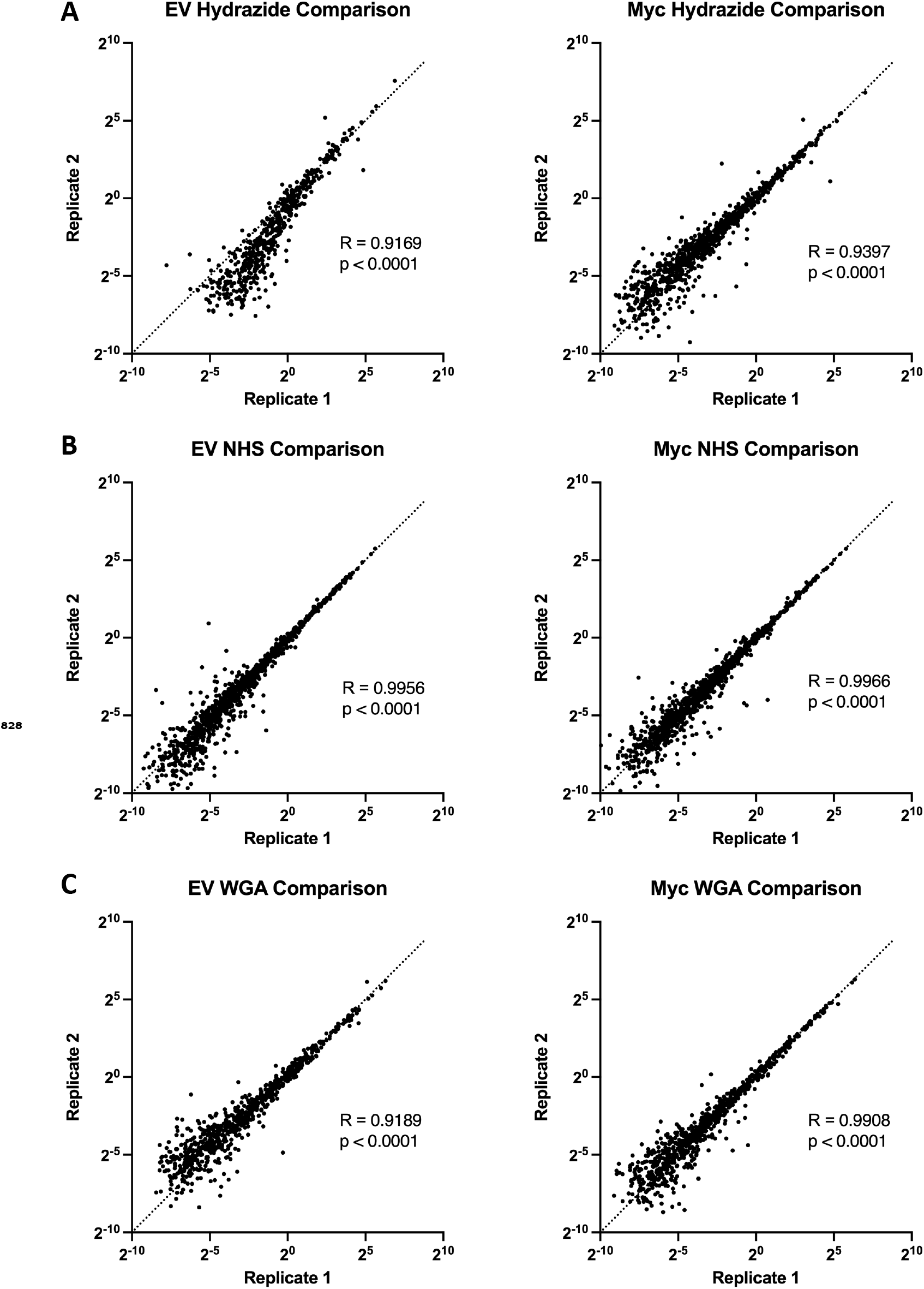
Comparison of replicates for different mass spectrometry methods show the WGA-HRP to have comparable reproducibility to Biotin-NHS or Hydrazide labeling. (A) Spearman correlations of TIC normalized data from replicates of Hydrazide EV and labeling. Myc cells. (C) Spearman correlations of TIC normalized data from replicates of WGA EV and Myc cells.

**Figure 4–Figure supplement 1.**
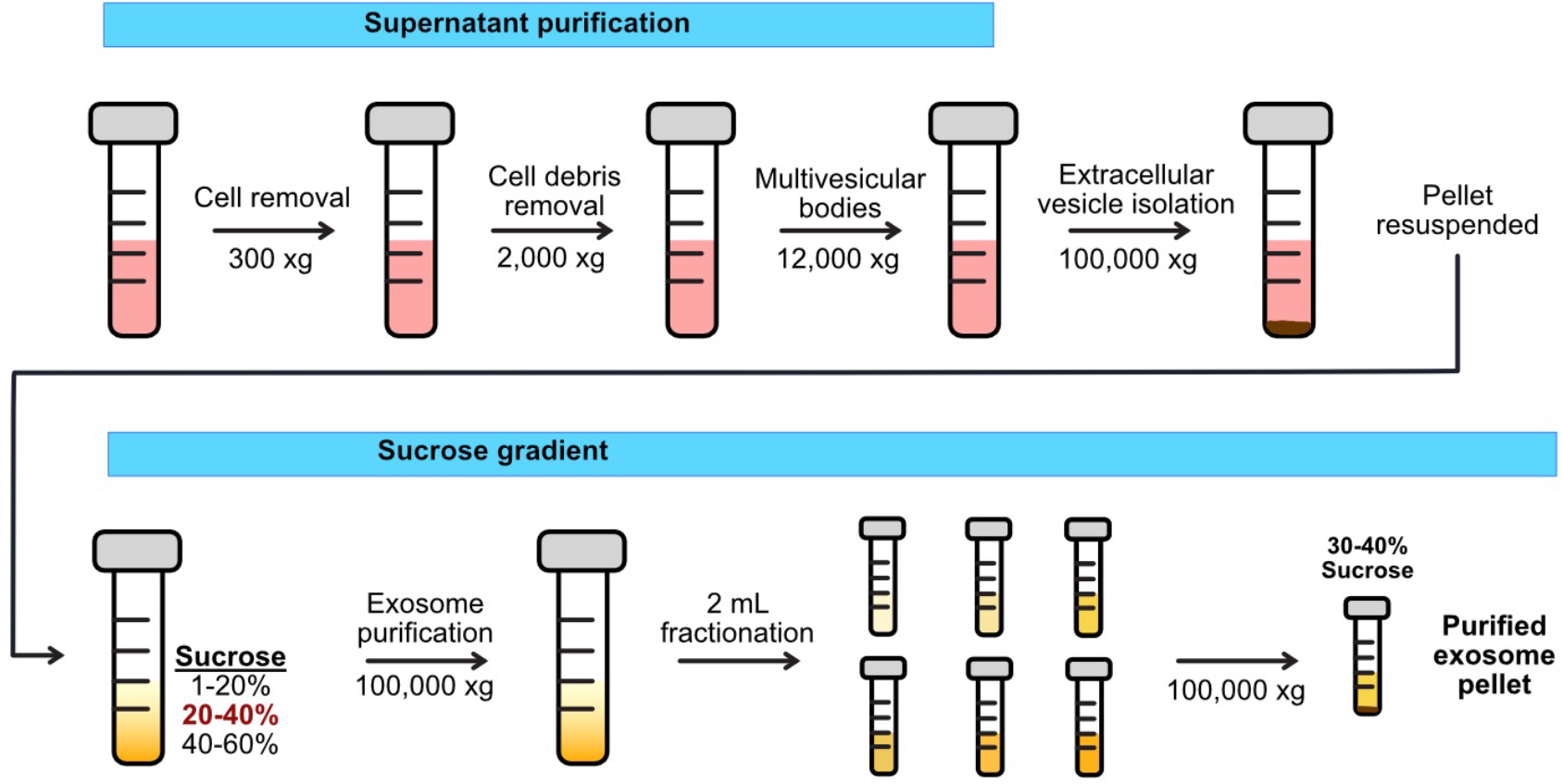
Workflow for exosome isolation from cultured cells. Media from cells undergoes serial centrifugation in order to isolate a mixed population of extracellular vesicles. Exosomes are isolated through sucrose gradient isolation and subsequent centrifugation.

**Figure 4–Figure supplement 2.**
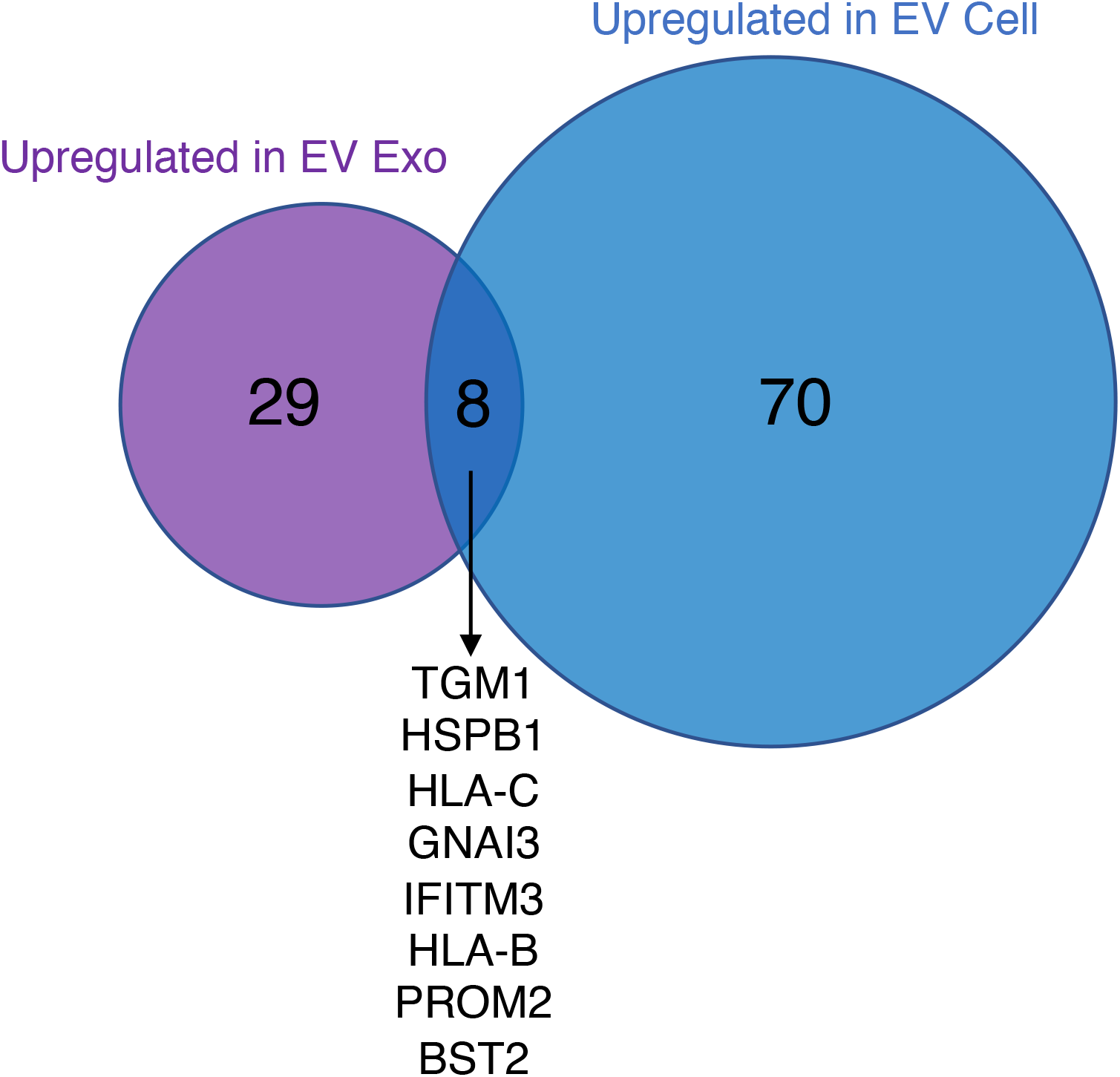
Venn diagram of enriched targets (>2-fold) in the EV Cells and EV Exosomes. Targets that were found enriched in the EV Exosomes compared to Myc Exosomes (purple) and the EV Cell compared to the Myc Cell (blue) were compared. The eight overlapping enriched targets are common between EV Cell and EV Exosome are listed in the center. the EV Exosomes compared to Myc Exosomes (purple) and th compared to the Myc Cell (blue) were compared. The eight ove enriched targets in common between EV Cell and EV Exosome a in the center.

**Figure 4–Figure supplement 3.**
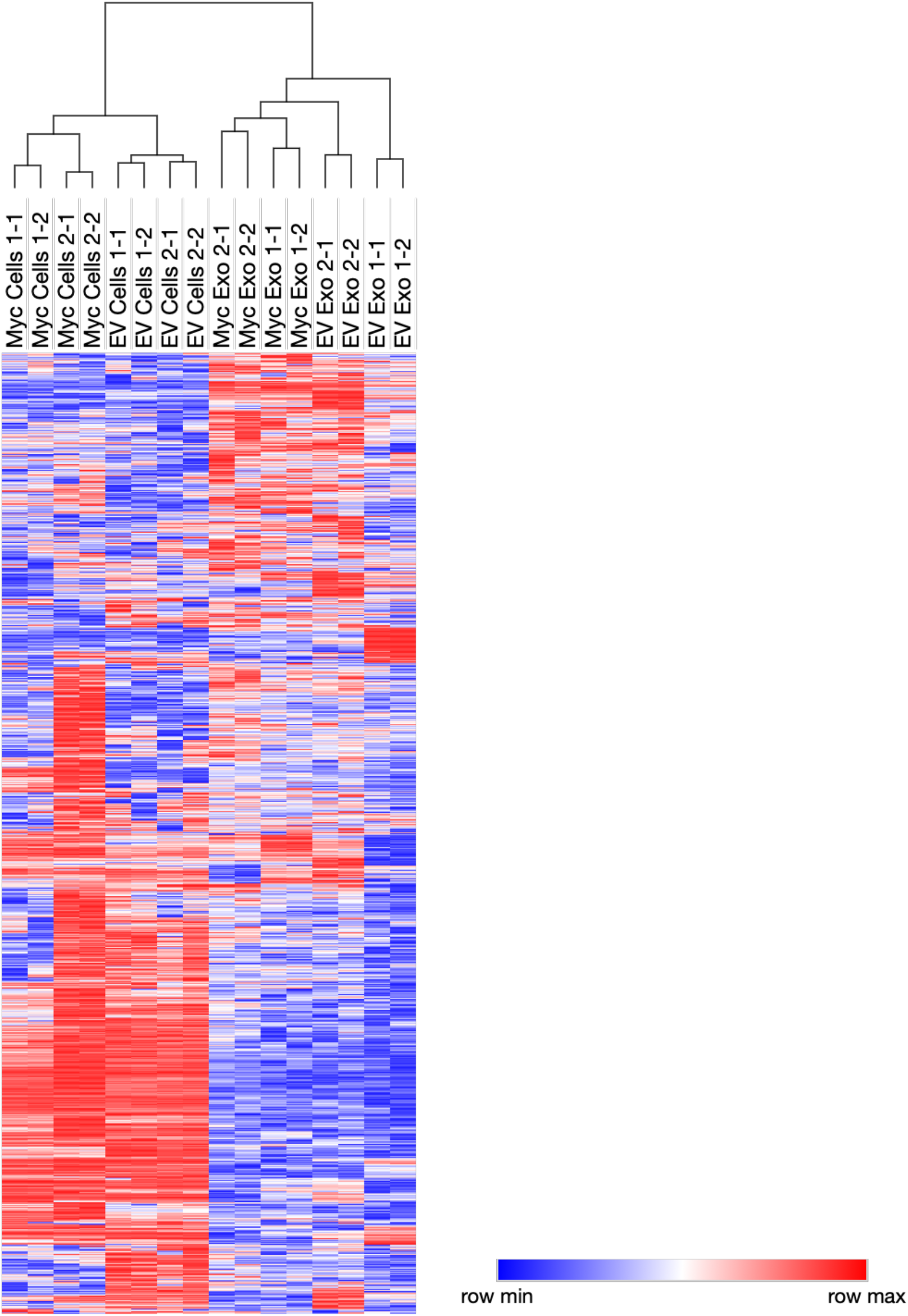
Heatmap comparison of biological and technical replicates of RWPE-1 EV/Myc cells and exosomes. Biological and technical replicates cluster together based on both oncogene status and compartment for exosome or cell surface. Proteins with no area values were assigned an imputed value using Perseus. Heatmap clustering is based off of the Pearson correlation between all replicates on both columns and rows. Heatmap was produced using Morpheus, https://software.broadinstitute.org/Morpheus.

